# Naturalistic paradigm reveals multi-component emotion dynamics in theta and beta bands using DENS dataset

**DOI:** 10.1101/2021.08.05.455187

**Authors:** Sudhakar Mishra, Mohammad Asif, Uma Shanker Tiwary

**Author notes:** Corresponding author, (S. Mishra).

## Abstract

The emotion research with artificial stimuli does not represent the dynamic processing of emotions in real-life situations. The lack of data on emotion with the ecologically valid naturalistic paradigm hinders the knowledge of emotion mechanism in a real-world interaction. To this aim, we collected the emotional multimedia clips, validated them with the university students, recorded the neuro-physiological activities and self-assessment ratings for these stimuli. Participants localized their emotional feelings (in time) and were free to choose the best emotion for describing their feelings with minimum distractions and cognitive load. The obtained electrophysiological and self-assessment responses were analyzed with functional connectivity, machine learning and source localization techniques. We observed that the connectivity patterns in the theta and beta band could differentiate emotions better. Using machine learning, we observed that the classification of affective self-assessment features, namely dominance, familiarity, and self-relevance, involves midline brain regions responsible for mentalization and event construction activity compared to valence and arousal, which were mainly associated with lateral brain regions. This finding advocates the need for more than two dimensions for emotion representation. In addition, the channels with high predictability were source localized to the brain regions in default-mode, sensorimotor and salience networks. Hence, in this naturalistic study, we find that the domain-general systems contribute to emotion construction.

## 1. Introduction

The controlled studies lack in capturing the complicated dynamics of real-life emotional events. These traditional studies in affective neuroscience often rely on varying one aspect of a stimulus at the time and use stimuli such as facial expressions, very short animated movies, and static images(e.g., IAPS pictures)[64]. However, theories have reported that emotional experience is a multi-component phenomenon and it emerges out of the integration of information from these components[69, 66]. The naturalistic stimuli, with their contextual and narrative structure, provide the possibility to consider many components simultaneously and induce reliable and ecologically representative neural responses during emotion stimulation [31, 4, 2]. In addition, they preserve the natural timing relations between the constituting functional components and the rapidly changing nature of emotions. The functional components of emotions range from low-level sensory and stimulus feature level to abstract high-level where observer’s evaluation and emotional category representations, integrating different processes including perception, memory, prediction, language, and interoception, are encoded [12, 67, 42]. These features across the processing hierarchy are necessary for a comprehensive model of emotional processing [12].

However, the emotion features related to the observer’s evaluation are averaged for the whole duration of naturalistic stimuli [64], which is not representative of the varying nature of emotions. For instance, different emotional categories can be felt at different times instance while watching the naturalistic stimuli. Hence, with a naturalistic paradigm, emotion features should be extracted continuously. In addition, averaging distills sensitivity of the emotion features in subtle neurological responses and confines them to dominated responses in sensory areas [64]. On the contrary, dynamic tracking of neural correlates of emotion features reveals the subtle activity in constitutive large-scale brain networks [57, 58].

However, dynamically collecting the emotional features (especially self-reports) imposes some challenges [68, 41]. For instance, collecting the self-reports during the experiment dynamically can alter the experience of emotion itself. On the other hand, the retrospective collection depends on autobiographical memory and can raise biases across subjects depending on their capability to recall [6]. Also, repetitive viewing effects can bias the ratings and underlying neural effects [6]. Hence, an experiment paradigm, which can record the participants feedback dynamically, with minimal distraction during emotion processing and minimization of the memory recall bias-ness, is needed.

The conceptualization of emotion as a dimensional construct advocates the need for more than two dimensions to describe emotional experience [20]. Generally, in literature other than valence and arousal, dominance is proposed as the third dimension [40, 80]. However, consideration of dominance as the dimension is still debatable as it rarely explains more than 15% of the variance in subjective rating [40]. The circumplex model of affect [14] acknowledged that to define a prototypical emotional episode including core affect dimensions (viz. valence and arousal), other components like attribution of cause and meta-cognitive judgement are needed. For example, experientially, fear and anger are distinct emotional states. However, the two-dimensional model will project it as a high-arousal and low valence state, which does not reflect the subjective experience of these emotions. In addition, the perception of emotion is not static across time or situations. This dynamicity in the perception of emotion is introduced by subjective meta-cognitive evaluation of the situation at a particular time [19]. While researchers endorse the need for more than two emotional dimensions, no consensus exists about what it could be.

In addition, the role of domain-general systems such as interoceptive (or salience network) [24], and DMN in emotion processing is also debatable, majorly due to research with artificially controlled stimuli on understanding the emotion mechanism, which generally covers the narrow aspect of this abstract phenomena. For instance, primarily, the emotion research reported activity in the amygdala, as evident from the emotion activation related meta-analysis image (fig-1 created using neurosynth meta-analysis python API [88]). However, the functional activations of emotional experiences do not reside in a single region or network (e.g., amygdala or limbic network) but are distributed across multiple, large-scale functional networks [67]. Unfortunately, investigation in emotion research has explored these large-scale networks to a limited extent, such as DMN contains some nodes which are affiliated with affect processing (e.g., vmPFC in the representation of pleasant and unpleasant states) [83]. Given the multi-functional roles attributed to these domain-general networks [60], it is mainly unclear that what constitutive roles they might play in the processing of emotions.

**Figure 1:**
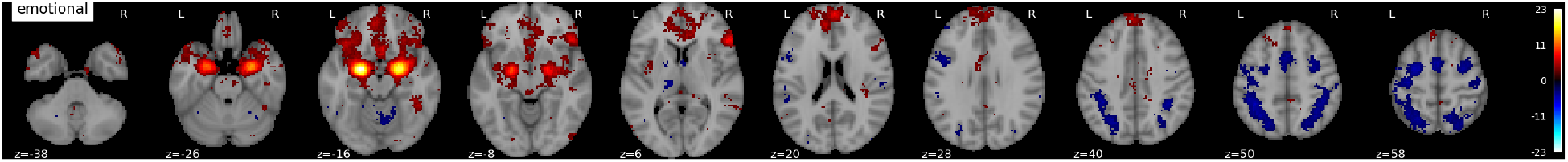
The statistical map for emotion activity is created from meta-analysis api provided by neuroSynth https://neurosynth.org/

Moreover, primarily, the extensive literature on emotion research uses the limited component of emotional features, e.g., voices, emotional pictures, facial expressions. The brain activity related to these features is mainly reported in high-frequency bands. For instance, a recent survey [34] shows that the gamma band is correlated with the processing of emotions and emotion memory. However, the constructionist point of view of emotion [12] posits that emotion is the constructed perception that involves brain waves supporting endogenous mental event construction and active prediction [15]. Hence, the probe on the spatio-temporal dynamics of emotion, situated in a context, can reveal the brain waves, generally, reported in the construction and perception processes [15, 76].

To address the issues mentioned above, we asked the following questions. 1) Can the naturalistic paradigm reveal the processing dynamics of emotion, which is not reported with the controlled experiments using artificial stimuli?; 2) do emotion representation require more than two dimensions (viz. valence and arousal)?; 3)Neurologically and functionally, what are the missing emotional components which are essential for the emergence of emotion and its experience?

To address the questions described above, we have designed an EEG experiment with an ecologically valid experimental setting (described in the section 2.3 and data article [52]). The raw signal is preprocessed and analyzed with five different kinds of analysis. First, we analyzed the relatedness and monotonicity of self-assessment dimensions. Second, we divided the EEG signals for each emotion click into eight segments and then calculated the functional connectivity pattern for each emotion group, frequency band and segment. Third, we calculated how much information each channel has to predict the self-assessment dimensions using machine learning. Fourth, we source localized these channels to obtain the brain regions. Finally, the correlations between power for each EEG channel and the self-assessment dimensions are calculated to understand the association dynamics of brain physiology with emotional ratings.

We found that emotion representation needs more than the traditional valence and arousal dimensions to reveal the construction dynamics. In addition, these construction dynamics are processed by domain-general systems such as dmn, sensorimotor and interoceptive (or salience network) in theta and beta bands. The rest of the article describes the experiment and methodology, followed by the results and discussion. We conclude with some suggestions for future experiments.

## 2. Material & methods

### 2.1. Stimuli Collection, Selection, and Validation

The procedure for stimuli collection, selection and validation is depicted in fig-2a (also see supplementary fig-1). For 372 affective terms, approx. 15000 multimedia videos with 150 comments per video were collected using youtube API. Textual analysis of these comments is done to select the videos with less ambiguity on positive and negative affective feelings. For the videos with high polarity, the multimedia content analysis (audio-visual features explained in data article [52]) is done to find out the three 1-minute excerpts from each video which are represented on the periphery of valence-arousal space (shown in the supplementary fig-2). This way, 220 stimuli were collected for the validation study with 666 participants. The stimuli which were seen at least by 35 participants and had the value of the SD less than three for valence, arousal and dominance were selected from the validation study. The intermediate result of valence and arousal ratings, during validation with 69 stimuli, is shown in fig-2b. We finally selected only 15 stimuli from the validation study with the most consistent responses for our EEG experiment.

**Figure 2:**
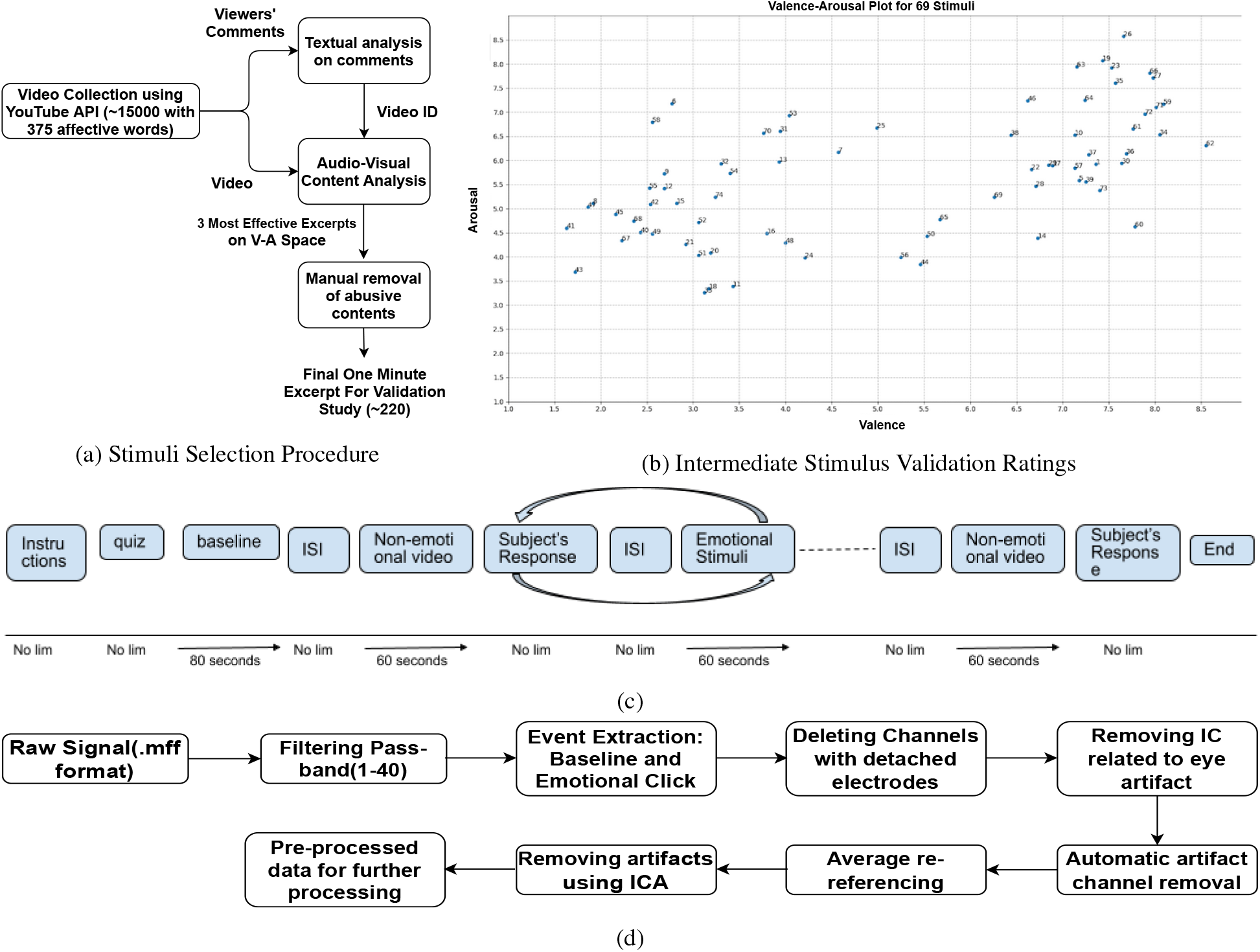
Experiment: (a) Stimuli selection procedure; (b) Scatter plot of subject ratings, collected during stimuli validation, on valence-arousal space; (c) EEG experiment paradigm; (d) Pre-processing pipeline.

### 2.2. Participants

Participants between 18-30 years of age were recruited. Inclusion criteria for all participants were based on Axis I assessment defining the criteria for mental health [9] (participants’ self-ratings submitted online. Also, see supplementary fig-6). We made sure that the self-rated scores are less than the threshold criteria of further enquiry as mentioned in the DSM-V manual [9]. The Institutional Review Board approved this research at the Indian Institute of Information Technology, Allahabad. All participants provided written informed consent and were compensated for their time.

Concerning participant characteristics, gender distribution was 85% males and 15% females. The average age of participants was 23.3 ± 1.4 years. Participants reported a time-difference between awareness about emotional feeling and mouse click response (described in section 2.3) which were 2-3 seconds (fig-3c) [32]. In addition, participants’ overall cognitive load, interference due to click task in emotion processing and general mood during the experiment are self-rated (in supplementary fig-6). Overall the participants’ cognitive load were towards the lower side of the scale. The average mood rating was 6.73 ± 1.97 on 1(unpleasant)-10(pleasant) scale, and their emotional feelings were least interfered by the clicking task.

**Figure 3:**
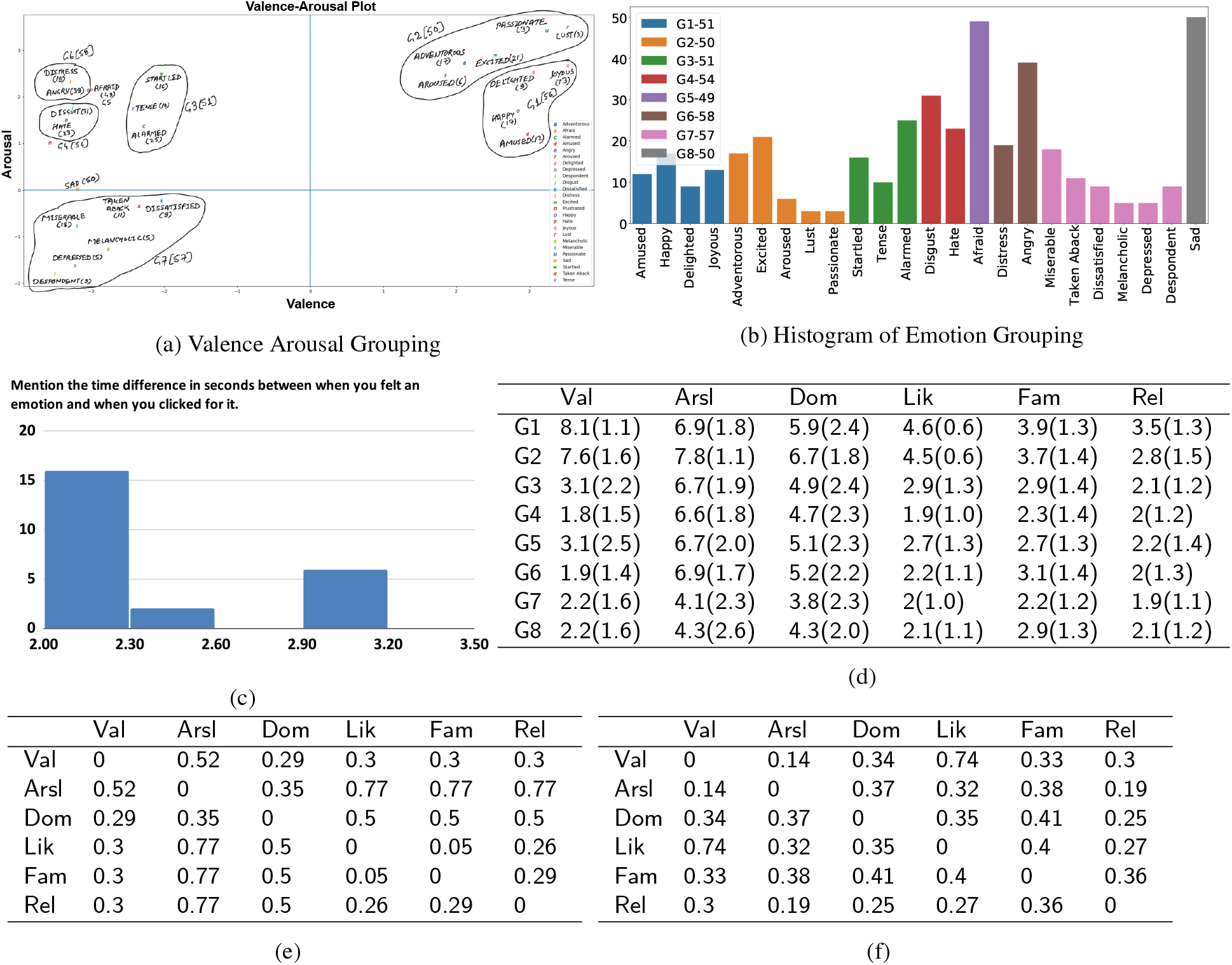
Behavioural Analysis: (a) Grouping of emotions. (b) Histogram depicting the count of individual emotions and their assignment to different emotion groups. (c) The difference in time of awareness about emotional feelings and mouse click. (d) The mean(sd) is tabulated for emotion-wise ratings by the participants. (e) Result of K-S test with *p* – *value* ~ 0.05. The values here represent the statistic D, the maximum absolute difference between the two cumulative distribution functions. The higher and lower differences would imply higher and lower dissimilarity of the two distributions. (f) Spearman rank-order correlation coefficient, a non-parametric measure, is performed to test the monotonicity of the relationship between two samples with the *p* – *value* ~ 0.05.

### 2.3. Experiment Paradigm

We have designed an EEG experiment along with ECG and EMG recordings (as shown in fig-2c). First, participants are provided with a training session with four one-minute multimedia stimuli. During training, participants are informed about the experiment protocol, taken a quiz (to assure they understood the protocol well), got familiar with the procedure and are trained to mark the moment with the mouse click when feeling any emotion (for the detailed procedure, please see data article [52]).

The baseline signal (eyes open looking at the cross mark) for 80 seconds is recorded. Then, 11 one-minute multi-media stimuli (two non-emotional and nine emotional) are presented. Every stimulus is preceded by an inter-stimulus interval and followed by response windows to respond on self-assessment scales: valence, arousal, dominance, liking, familiarity, relevance and emotional category selection. While watching the stimulus, participants marked the moment of emotional feeling with a mouse click. They can click any number of times. While selecting an emotion category for these clicks, the participants are given a list of emotions in all four quadrants of valence-arousal space in a drop-down menu so that participants can freely select the emotion they felt rather than imposing the emotions on them as done in previous studies[45]. To help recall the felt emotion, participants were shown the frames extracted around the time duration when they did mouse click for having felt any emotion. With this simple trick, we localized a participant’s emotion temporally with less distraction and cognitive load. The collected electrophysiological data (EEG and ECG) is preprocessed in Matlab scripts with the help of EEGlab functions.

### 2.4. EEG Data Acquisition

Participants sat in the immovable chair with an armrest and facing the screen kept at a 1-meter distance from the participants. The stimuli were presented on a 15.6-inch screen display (1024×768 resolution) using a dedicated computer system with Intel Core i5 3rd generation, 3.2GHz. However, the resolution of the video stimuli was reduced to 800×600. For the audio, Sennheiser CX 180 Street II in-Ear Headphone was used. Participants were allowed to adjust the audio volume to their comfort level during training. For response, a mouse and a keyboard were also placed.

For recording EEG, 128 Channel Geodesic EEG System 400 was used. 128 EEG channels cap (HCGSN v.1.0) and three peripheral signals (one ECG and two EMG electrode pairs with Physio16 Package) were correctly placed. The raw physiological signals were recorded at a sampling rate of 250 Hz using net station software. A dedicated computer system, iMac, was used for recording and storing the raw signals.

### 2.5. Pre-processing

The raw data is available at [51]. The preprocessing steps are performed using eeglab functions (shown in fig-2d). The raw signal is imported, which is already referenced to Cz(reference electrode, EGI default). However, we performed average re-referencing during the preprocessing. The raw EEG signal is filtered using a fifth-order Butterworth bandpass filter with the low-cutoff 1 Hz and high-cutoff 40 Hz. From the filtered signal, the event segments corresponding to baseline state, pre-stimulus, and the click are extracted with time-duration 10s to 70s, −3s to 0s, and −6s to 1s, respectively. The extracted duration for the click event is from 6s before the click event to 1s after the click.

Next, the concatenated signals were manually checked. The channels and samples with very high amplitude, probably caused by electrodes detachment, are removed manually (The manual rejection sheet is online). Then, we applied the ICA to remove the eye blink artefact so that high amplitudes of eye artefacts should not misguide our next step, which is automatic channel rejection. Then, the second ICA is applied for any other remaining artefacts in the signal (including eye, muscle, heart, line, and channel artefacts). We kept independent components with the probability of brain activity more than 0.3 (adding the probability factors for all the artefacts, including brain signal, equal to 1). After completing the preprocessing steps, the individual events corresponding to baseline state and emotion clicks (separately) are stored in .mat files for further data analysis.

For ECG, the channel data is extracted and bandpass filtered with low-cutoff 0.6Hz and high-cutoff 40Hz (all other filter parameters are the same as with the EEG processing). Only segments corresponding to the time duration for the resting state event and click event (duration mentioned in the above paragraphs) were considered from the continuous signal. The ECG signals are shown in supplementary fig-7.

### 2.6. EEG Analysis

#### 2.6.1. Emotion Grouping

We have grouped the emotions based on two criteria. First, the distance-based proximity of mean (calculated for each emotion) on V-A space, and second, approximately equal samples across emotion groups. Based on these criteria, we obtained the emotion groupings (as shown in the fig-3a & 3b). The mean (sd) ratings of self-assessment scales for each group are presented (tab-3d). We utilized this grouping for further analysis.

#### 2.6.2. EEG Signal Segmentation

The duration of extracted EEG signal corresponding to each emotional event is 7 seconds. The sampling rate was 250Hz. Each emotional event signal is divided into segments containing 250 samples and 75 samples overlapping with the previous and next segment. In this way, we got eight segments (without zero-padding) to perform the spatiotemporal dynamic analysis.

#### 2.6.3. EEG Dynamic Functional Connectivity

We used phase-locking value (PLV) as a measure for calculating connectivity between EEG Electrode pairs. For PLV calculation, the real signal is transformed into an analytic signal using the Hilbert transform. The instantaneous phase is then calculated. The instantaneous phase difference between pairs of electrodes is quantified as PLV value. For each frequency band, segment, and emotion group, PLV values are calculated among pairs of electrodes. Then the connectivity matrix for each emotion group is contrasted with the baseline state connectivity matrix, and a non-parametric approximate permutation test (corrected for multiple hypothesis comparisons using single threshold technique) is calculated (see section 2.6.6).

##### Calculating distance between connectivity patterns

The Euclidean distances between the connectivity vectors for all emotion pairs are calculated. Henceforth, we obtained connectivity distance for each emotion group pair (for all frequency bands and segments). We utilized these connectivity distances to find out which frequency bands have relatively more information. To this aim, we calculated the median value of connectivity distances among pairs of emotion groups for each frequency band across the segments. These median values are annotated in the histogram plot of fig-4a. Then, the frequency bands with the median value above the global threshold (*G_cst_*=median of connectivity distances calculated across all emotion pairs, frequency bands and segments) are reported as the frequency with relatively more information.

**Figure 4:**
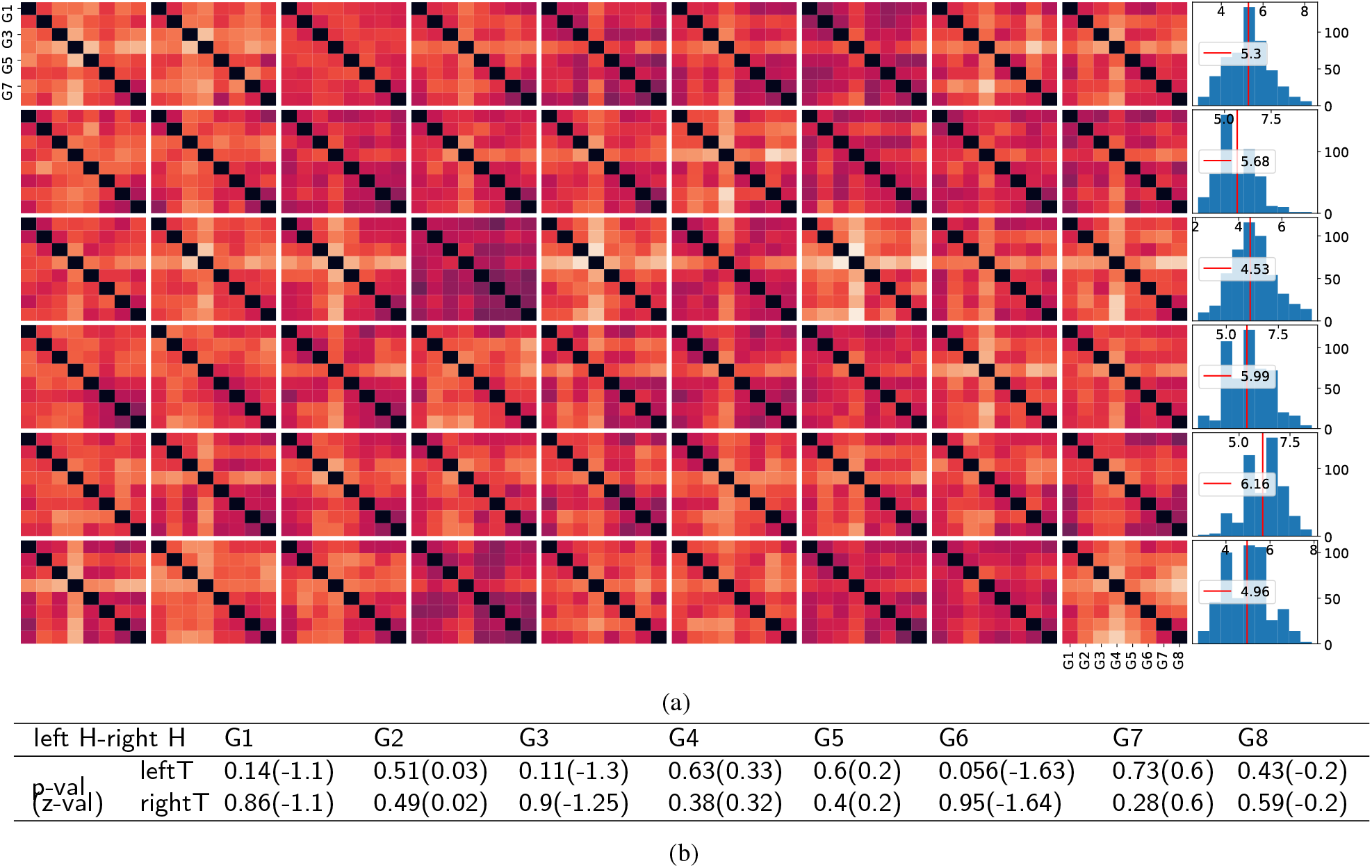
The euclidean distance between connectivity patterns for different emotion groups: (a)The Euclidean distance is calculated among pair of emotion groups and plotted as a heat map for each frequency band (row-wise: delta, theta, alpha, lower beta, upper beta, and gamma) and each segment (column-wise: seg-1 to seg-9). Black represents the lowest, and white represents the highest distance. A more distance is better. The last column is the summarizing histogram with the annotated median statistic. We calculated the global threshold (*G_cst_* = 5.44) by taking all the distances into account. Only theta and beta band has more values of representative normalized distance among connectivity patterns of emotion conditions. (b) From the EEG Connectivity matrix, nodes on the left and right hemispheres are collected (only nodes making significant links calculated in connectivity estimation among EEG electrodes). For each emotion category, the active left and right electrodes were counted. These values represent how many electrodes from the left and right hemispheres have participated significantly in the connectivity. We then performed Wilcoxon signed-rank test for each emotion category with the contrast between electrodes on the left and right hemispheres. The p-value and z-value are tabulated. Except for group-6, no other emotion groups show hemispheric specific significant activation. The supplementary connectivity diagram for each emotion condition is shown in (supplementary fig-3).

#### 2.6.4. EEG Features Calculation and k-NN classification

To perform the classification analysis, we calculated 20 representative features which include Detrended fluctuation analysis(DFA), Power Spectral Intensity(PSI), Relative Intensity Ratio(RIR), Petrosian Fractal Dimension(PFD), Higuchi Fractal Dimension (HFD), Hjorth mobility and complexity, spectral entropy, SVD Entropy, approximate entropy, fisher information, and Hurst exponent [11] (see supplementary section-SI: EEG Features).

In the classification analysis, we quantified the contribution of information of different scalp sites (recorded using EEG electrodes) in predicting the ratings of the self-assessment scales. To this aim, we calculated the predictive power of each EEG channel using the k-NN classification algorithm. For each electrode, a vector of the features (20 features) mentioned above is calculated. Then, we considered each electrode and trained kNN classifier with 70% data to classify two classes per scale (high & low) created by thresholding valence, arousal, dominance, liking, familiarity, and relevance, respectively, with the values 5, 5, 5, 2.5, 2.5, 2.5. The k-NN classifier is trained to learn an optimal boundary that can perform the binary classification with the least error. The remaining 30% data is used for testing the trained k-NN classifier and predicting the classes. The accuracy parameter is used to assess the performance of the classifier. The reported accuracy is the mean accuracy for 20 classification trials with sub-sampling from the original number of samples for each scale. The prediction accuracy is then plotted on the topo plots (fig-5b).

**Figure 5:**
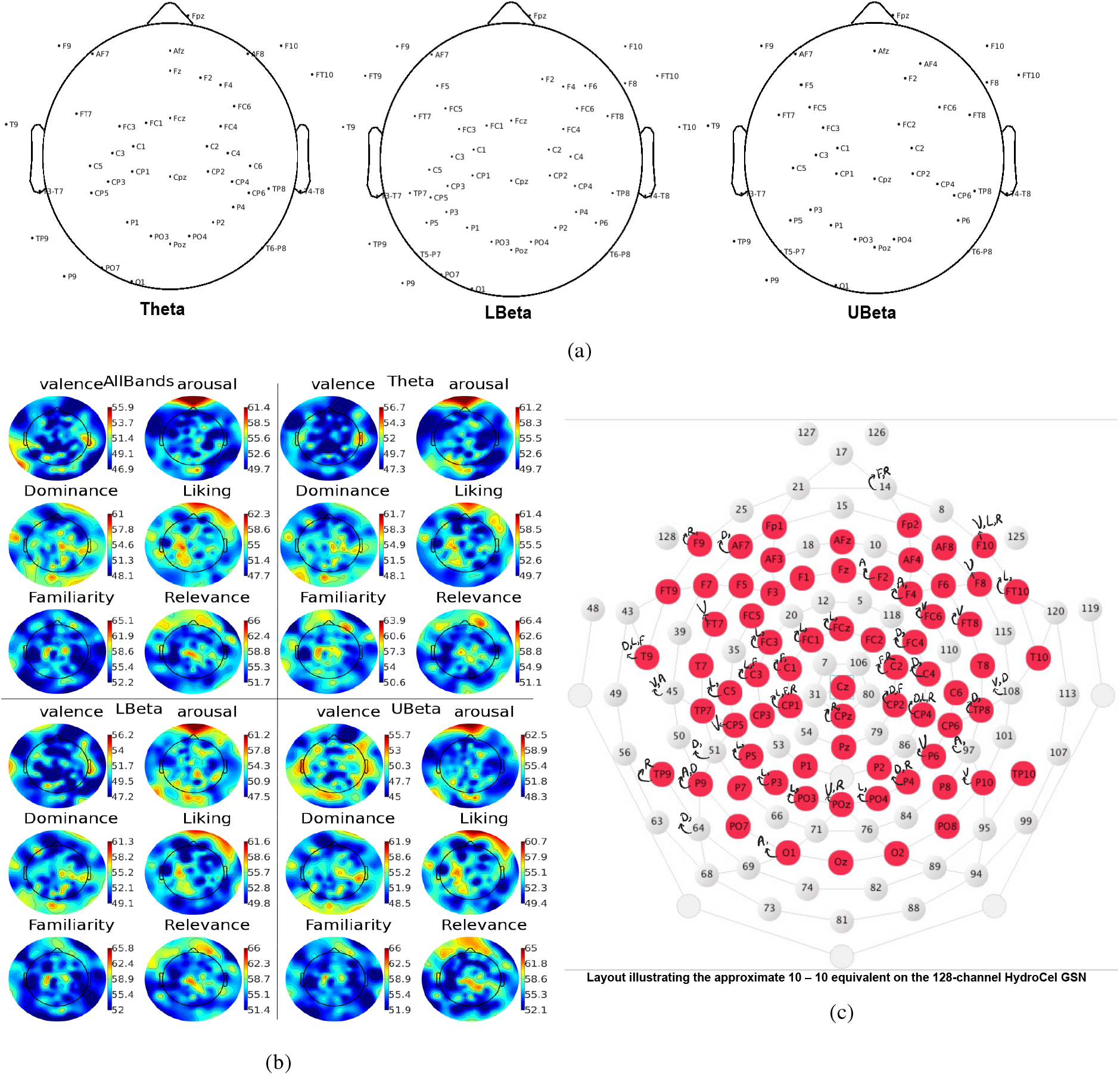
Channels with high prediction accuracy for self-assessment scales: The plotted results are obtained after the k-NN classification for each channel and retaining only those channels which are above the threshold (*G_pt_*). (a) Channels with high prediction accuracy are plotted in scalp layout for theta, lower beta and upper beta bands as connectivity patterns in these bands are more distinct. (b) The prediction accuracy for channels predicting self-assessment scales is plotted in the topoplot (plots for other frequencies are in supplementary fig-4). All Bands: signal containing 1-40Hz. (c) Annotated channel layout. Channels with more than the threshold are annotated with the corresponding self-assessment scale. Abbreviations: Valence(V), Arousal(A), Dominance(D), Liking(L), Familiarity(F), Relevance(R), Anterior Frontal(AF), Frontopolar(Fp), Frontal(F), Frontotemporal(FT), Frontocentral(FC), Temporal(T), Temporoparietal(TP), C(Central), Centroparietal(CP), Parietal(P), Parietooccipital(PO), Occipital(O).

#### 2.6.5. Extracting brain regions for selected electrodes using equivalent dipole source localization

As mentioned in the section 2.6.4, using k-NN classification we have obtained prediction accuracy for all the channels. Then, we calculated the global threshold *G_pt_* to select channels with high prediction accuracy using the formula

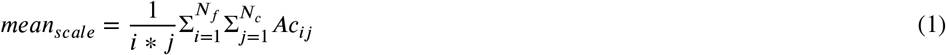

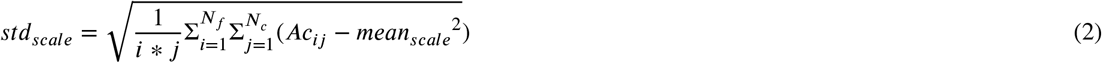

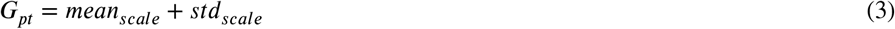

Where, Ac:Accuracy, *N_F_*:Frequency Bands, *N_c_*:Channels. Channels with accuracy higher than *G_pt_* (From now on referred to as prediction channels) are shown in the annotated fig-5c. The channels are co-registered to the matching electrode locations template (10-5 electrode system) associated with the BEM volume conduction head model (supplied with EEGLab). The transformation matrix for co-registration is [-0.2667, −18.3138, −11.2267, 0.0282, −0.0104, −1.5501, 10.1523, 10.2578, 10.6686]. The prediction channels are source localized to the dipoles with 40% residual variance threshold. The resultant coordinates for these dipoles are in Talairach coordinate system. These coordinates are then converted to MNI coordinate system. The anatomical annotation of the MNI coordinates is performed using the “Talairach Client - Version 2.4.3” application. The obtained brain regions are shown in the histogram (fig-6b), and brain regions higher than the median value are shown in fig-6a

**Figure 6:**
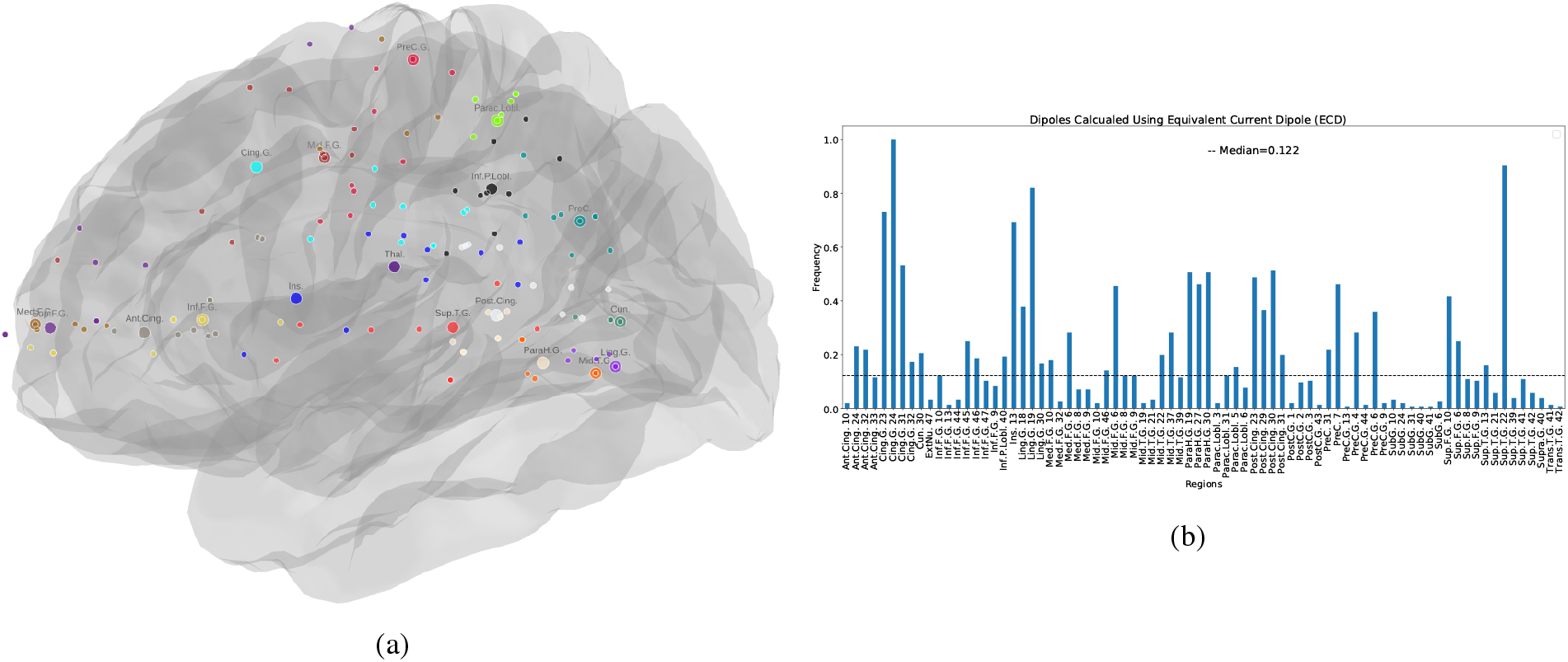
The Equivalence Current Dipoles. We have calculated equivalent current dipoles corresponding to independent components calculated for prediction channels for self-assessment scales. (a) Localized anatomical regions annotated on the brain. (b) The histogram shows the frequency of occurrence of the anatomical regions. The horizontal black line represents the median which is considered as the threshold. Regions above this threshold are depicted in (a). Abbr: inferior(inf), medial(med), middle(mid), cingulate(Cing.), Frontal(F), Gyrus(G), Insula(Ins), Anterior(Ant), Thalamus(Thal), Precentral gyrus(PreC.G.), Paracentral(Parac), Lobule(Lobl), Parietal(P), Precuneus(PreC), Cuneus(Cun), Posterior(Post), Superior(Sup), Temporal(T), Parahippocampal(ParaH), Lingual(Ling).

Fig-5c shows that self-assessment dimensions other than valence and arousal show higher accuracy around scalp sites, including frontocentral, midline structures, centroparietal and parietal regions. These regions are part of domain-general systems and are involved in event construction activity. It also explains why earlier research on emotions with only valence and arousal got a limited understanding of the mechanism of emotion.

#### 2.6.6. Statistical Significance Testing

##### Kolmogrov-smrinov test (K-S test)

The two-sample K-S test calculating the maximum absolute distance (D) between the cumulative distribution function (CDF) of two samples is performed. The null hypothesis is that two related or repeated samples have identical CDFs (for *p* < 0.05; tab-3e). The difference between CDF ranges from 0 to 1. 0 means two distributions are the same, whereas 1 means they are entirely different.

##### Non-parametric Spearman’s rank-order correlation

The Spearman rank-order correlation coefficient is a nonparametric measure of the strength and direction of association that exists between two variables. We performed the correlation test between each pair of self-assessment dimensions with the null hypothesis that two dimensions have the monotonicity of the relationship (for *p* < 0.05;tab-3f).

For each segment and frequency band, the spearman correlation test between the power of EEG channels and the self-assessment ratings-valence, arousal, dominance, liking, familiarity, and relevance was calculated. Only the significant correlation coefficients with *p* < 0.05 is plotted on the topoplot (fig-7).

**Figure 7:**
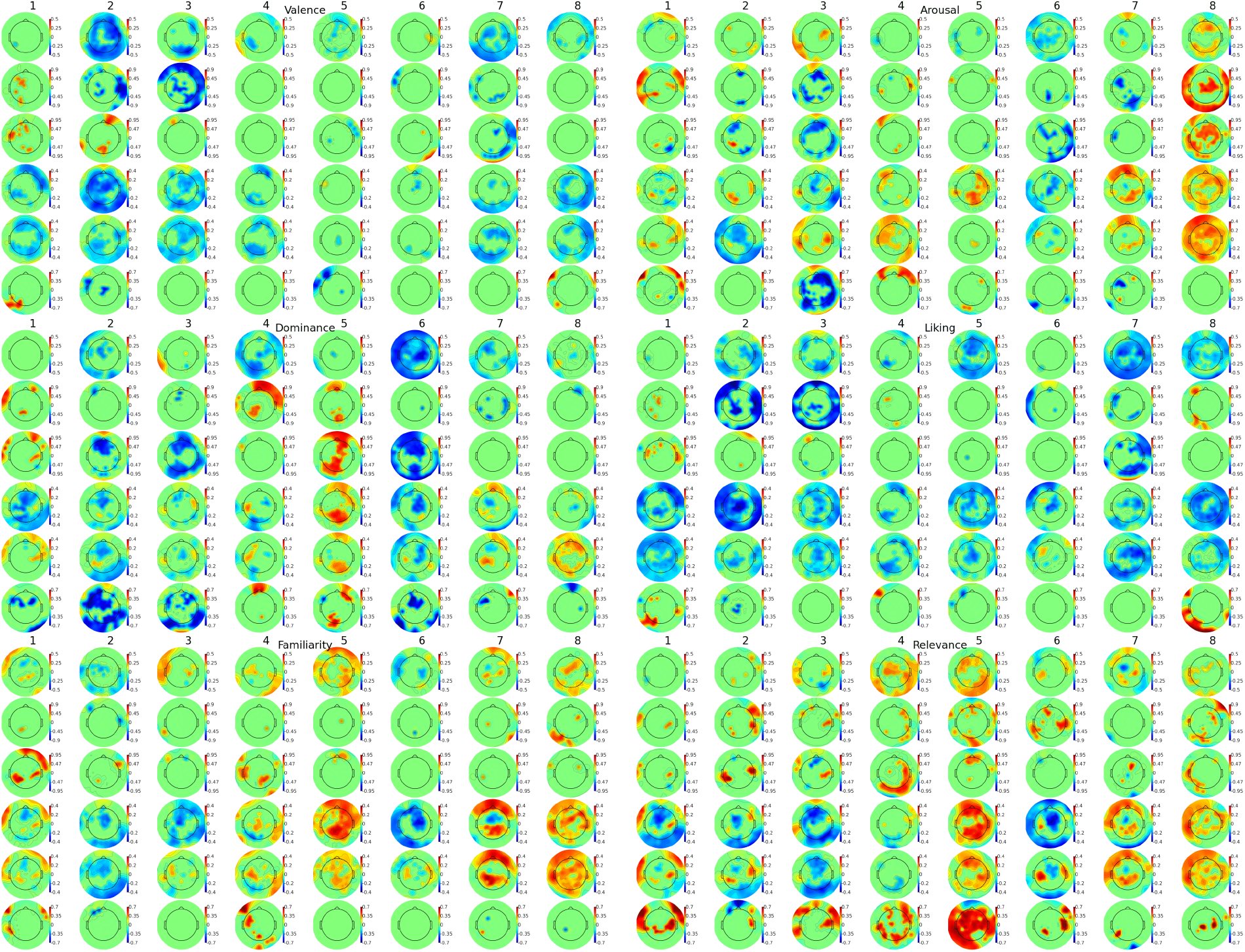
Dynamic correlation of prediction channels with self-assessment ratings: The colour coding in the topoplot, from blue to red, represents negative to positive correlation statistics with the statistical significance(*p* < 0.05). The spearman correlation is calculated between normalized power ratio (normalized with the resting-state condition) for each channel and subjective ratings for valence, arousal, dominance, liking, familiarity and relevance. This calculation is done segment-wise so that the correlation dynamics can be traced.

##### Non-parametric permutation test for functional connectivity

We performed a multiple-comparison non-parametric permutation hypothesis test for related data corrected for family-wise type-1 error(adjusted p-value ~ 0.05) to find the significant functional connection among pair of electrodes. The null hypothesis is that permuting the labels of related samples assigned to different experimental conditions leads to an equally likely statistic of interest. That means the statistic for actual labelling is in the confidence interval of permutation distribution obtained after a sufficient number of relabeling. Hence, the p-value is the proportion of the statistics greater than or equal to the statistics T corresponding to the correctly labelled data. This multiple comparison test is performed for all electrode pairs. The family-wise type-1 error rate, due to family-wise multiple comparisons, is controlled using a single threshold test. The key idea behind the single threshold test is that given a critical threshold (single threshold), functional connections with statistic values exceeding this threshold have their null hypothesis rejected. The critical threshold is decided using the distribution of the maximum statistics. This distribution of the maximal statistic is computed from the statistic image obtained after several permuted relabeling. The omnibus hypothesis at level *α* is rejected if the maximal statistic calculated for actual labelling is in the confidence interval 100*α*% of permutation distribution. The critical value is the (*c* + 1)^*th*^ largest member of the permutation distribution, where *c* = [*αN*]. If the statistic image passes the omnibus test, further, the voxel/electrode can be rejected if exceeding this threshold [1].

## 3. Results

We found that emotion is a multi-component phenomenon that needs more than two dimensions to capture its constitutive components and understand its dynamics. In addition, theta and beta brain waves facilitate emotion construction mechanisms by engaging brain regions forming dmn, sensorimotor and salience networks. Details are presented in the following subsections.

### 3.1. Behavioural Analysis

Using the k-s test, we observed significantly different CDF for all scale-pairs except for the pair liking-familiarity tab-3e. Hence, the self-assessment dimensions belong to different distributions (for *p* < 0.05) and having distinct information about emotion processing [65, 79]. However, liking and familiarity have the most negligible difference (0.05 with p-val=0.6) and confirm the null hypothesis that these two samples come from similar distributions. The distribution of valence ratings is maximally different from the distribution of arousal ratings (*D* = 0.52, *p* – *val* ~ 0). The distribution function for ratings of each scale is not entirely independent (as the difference between CDF is not 1) but have some relationship which we next probed with the correlation test.

Spearman rank-order correlation coefficient, a non-parametric measure, is performed to test the monotonicity of the relationship between two scales. We found that the dimensions are not independent of each other, and they share information with varying degrees of correlation (tab-3f). For instance, valence and liking are highly correlated (*r* = 0.74, *p* – *val* ~ 0) compared to valence and arousal (*r* = 0.14, *p* – *val* ~ 0), which are least correlated. Valence and arousal are significantly correlated with the dominance dimension (*valence* : *r* = 0.34, *p* – *val* ~ 0; *arousal* : *r* = 0.37, *p* – *val* ~ 0;). Whereas, relevance is least correlated with arousal(*r* = 0.19, *p* – *val* ~ 0). In a study by [61], participants are shown the least arousing images to elicit implicit self-relevance activity in them. In addition, familiarity is significantly correlated with other scales. In summary, the self-assessment dimensions are not significantly independent of each other and have varying degrees of relationships contrary to the k-s test (reported above). However, both tests agree on significant in-dependency of valence and arousal scales.

In addition, we observed that pairs of scales like valence-arousal and relevance-arousal are least correlated and might have a significant share of contribution in emotion processing[65, 35, 48]. With these results, we are commenting that valence, arousal, and relevance dimensions (*valence* – *arousal* : *r* = 0.14, *p* – *val* ~ 0; *valence* – *relevance* : *r* = 0.3, *p* – *val* ~ 0; *arousal* – *relevance* : *r* = 0.19, *p* – *val* ~ 0;) may be contributing more information (due to the lower degree of correlations among them) than valence and arousal with dominance or control dimension (*valence* – *arousal* : *r* = 0.14, *p* – *val* ~ 0; *valence* – *dominance* : *r* = 0.34, *p* – *val* ~ 0; *arousal* – *dominance* : *r* = 0.37, *p* – *val* ~ 0;).

We further tested the importance of dominance and relevance by calculating the euclidean distance between emotion pairs in valence-arousal-dominance(VAD) and valence-arousal-relevance (VAR) space. We found that the total number of pairs in VAR space with a higher euclidean distance than in VAD space was 280 compared to 272 pairs in VAD space. Furthermore, we performed a paired t-test for related samples and found that for the VAR-VAD contrast, the statistical score was 1.01 (with p-val=0.16). Although it is a weak significance, it supports the possibility that relevance is worth consideration in emotion research [65]. Maybe with more experiments and observations in future research, the possibility of consideration of relevance dimension gets more substantial support.

### 3.2. Theta and Beta Bands have differences in connectivity patterns of emotions

By calculating the difference between connectivity patterns (for the emotion groups), we found that connectivity patterns are more distinct in the upper beta band followed by lower beta and theta bands (fig-4). We did not consider other bands as they were below the global threshold (*G_cst_*, see section 2.6.3)). We also probed if emotion processing has lateralized functional connectivity. We did not observe functional connectivity lateralization for emotion processing (tab-4b).

### 3.3. DMN, sensorimotor and salience networks are localized for prediction channels

Considering the frequency bands calculated above, we used the classification and source-localization analysis to determine the scalp sites and brain regions that contain high predictability information in predicting self-assessment dimensions. Prediction channels for beta and theta bands are shown in fig-5a. Primarily, valence and arousal associated prediction channels are observed in lateral regions. On the other hand, for dominance, liking, familiarity, and relevance, prediction channels are mainly located near the midline brain regions (annotated in the fig-5c). The regions around the midline are mainly reported in mental construction and simulation of events [78]. Hence, considering only valence and arousal self-assessment dimensions may not reveal the mentalization and event construction component of emotion mechanism [12].

We source-localized EEG prediction channels by performing equivalent current dipole (ECD) source localization. We found source localized activity in the paralimbic systems, including cingulate gyrus (CG), insula (Ins), thalamus (Thal), parahippocampal gyrus (PHG), and inferior frontal gyrus (IFG). In addition, prediction channels are localized in the anterior frontal cortex, posterior temporoparietal junction(TPJ), somatosensory regions, medial parietal regions, pre-central motor regions, and visual cortex regions (as shown in fig-6a, 6b). The visual processing is localized in the cuneus and lingual gyrus. In the posterior medial section, the precuneus and cuneus are localized. The primary somatosensory (S1) and primary motor (M1) cortices occupy caudal and rostral portions of the paracentral lobule and are part of the sensorimotor network. The brain regions including CG/mPFC, IFG, PCC, IPL, TPJ, and PHG comprise the DMN network. Furthermore, the activity in the insular cortex and ACC forms the salience network.

### 3.4. Dynamic correlation of prediction channels with self-assessment scales in theta and beta bands

The valence rating is correlated in seg-2 with the prediction channels in the left hemisphere (viz. FT7, E45, CP5) in the lower beta band(fig-7). Whereas, during the seg-3, the correlated activity in the right hemisphere (in channels F10, F8, FC6, FT8, P6, P10, PO2 fig-7, fig-5c) is significant in theta (primarily) and lower and upper beta bands. During seg-2 and seg-3, we observed high power for low valence ratings and low power for high valence ratings. Similar results are observed in another analysis, on the same dataset, where the valence ratings were correlated with the power of different brain networks in seg-2, and seg-3 [53].

On the other hand, the correlation between arousal ratings and scalp electrode power for channels F2, F4, E97, E45, P9 is observed in seg-3 (fig-7). The correlation of arousal ratings with the prediction channels is primarily observed in seg-3 compared to seg-2 in valence. It might be that arousal activity is being modulated by the valence activity, which is also reported in an ERP study [30]. Although, further research is needed to understand the temporal dependence between scales.

During seg-6, prediction channels in the right central, centroparietal, parietal regions are negatively correlated with the dominance ratings in the beta band (fig-7). In addition, in seg-7, the correlated activity in the left parietal and parietooccipital regions in the beta band is observed (fig-7). These correlated activities associate control and regulated behaviour with participants’ subjective feedback. We speculate that the participant is regulating his/her emotional feelings to perform the goal-directed action (mouse click) [49].

For familiarity ratings, prediction channels correlated activity is observed in frontal pole, central and centroparietal regions during seg-5 in the beta bands (fig-7). A recent study explaining the familiarity of the context in a predictive coding framework shows that prediction error is the key to learning and moving from unfamiliarity to familiarity. In other words, if the prediction accuracy is high (equivalent to low prediction error), the familiarity of the content is inferred high [7]. Participants in our experiment are instructed to rate familiarity based on the prediction accuracy. Hence, the assessment of prediction accuracy might be delaying the correlated familiarity activity compared to valence and arousal. However, familiarity influences the valence and arousal activity which in turn depends on the prediction accuracy [36, 53].

The correlation of prediction channels for relevance is observed in the left centroparietal in the upper beta band (seg-2), right parietooccipital & centroparietal in the beta band (in seg-3 & seg-5) and right central regions in seg-6 (fig-7). Primarily, the relevance is correlated with the activity in the midline regions. In addition, compared to other scales, the correlation of relevance is observed in all segments. The self-relevance related autobiographical memory, which contains implicit knowledge about emotions, can be an automatic process and take part in the construction of emotion experience[37, 12].

## 4. Discussion

In this work, we adapted the naturalistic paradigm to understand the emotion processing dynamics. The naturalistic paradigm reveals the multi-component emotion processing in a natural, wild and dynamic environment, which lacks in the controlled experimental design using the artificial stimuli [64]. With the naturalistic paradigm, the continuous brain activity, representing the emotion dynamics emerging within the underlying context, can be recorded and analyzed to come up with the novel conclusions [64]. For instance, in our analysis, we observed that the dynamic connectivity pattern of emotions is distinctly presented in the theta and beta bands. In the literature, mostly, high-frequency bands are reported for emotion processing [34, 87]. The reason behind this is that the controlled stimuli consider a single component, e.g., facial expressions or body movements, or emotional pictures/scenes, which may be revealing the reaction to stimuli rather than the endogenous brain dynamics which is facilitating the interaction with the environment, thus the construction of the emotional episode.

In addition, it is reported that emotion is a constructed phenomenon [12]. The emotional episodes are reconstructed from experience, at the same time, accommodating the salient information from the current context [12]. The re-construction process of emotional episodes is not instantaneous but emerges from continuous interaction with the present situation. Hence, emotions are situated within the context [82], and cannot be captured with instantaneous reactions to pictures [64]. Our analysis found the default mode network for the re-construction process and visceromotor networks to embody the experience of reconstructed emotional episodes.

We have also observed that valence and arousal dimensions alone are inadequate in capturing the neural components essential to the re-construction of the emotional episodes. The neural correlates of valence and arousal are not distributed around the mid-line brain regions, which are reported as correlates of mental simulation and event construction [78]. These brain regions are associated with dominance, familiarity, and relevance dimensions. Hence, we report that consideration of more than two dimensions (viz. valence and arousal) is necessary to capture the neural dynamics of emotion construction. Moreover, the consideration of dominance, including valence and arousal in emotion representation, is also reported in a study from our lab [80]. On the contrary, we found that relevance has the least correlation with valence and arousal. Hence, the relevance can be considered as a representative self-assessment dimension of emotions [65]. The following sections discuss these findings.

### 4.1. Emotion Processing in Theta and Beta Bands

We observed the distinct connectivity patterns for different emotion groups and the correlation dynamics for self-assessment features-valence, intensity, dominance, liking, familiarity, and self-relevance-mainly in the beta and theta bands. In the literature, the beta band is reported in diverse functions, including maintenance of information in working memory, motor planning, content specific modulation, decision making, top-down perceptual processing, long-range communication, and preservation of the current brain state [76].

However, recent evidence suggests the role of the beta wave, particularly in content-specific reactivation and maintenance during endogenous information processing as demanded by the current task [76]. Furthermore, the maintenance of the activated information and integration with the information acquired in the current context is supported by the computational model of cell assemblies [46]. Beta-synchronized cell assemblies of superficial pyramidal cells are self-sustaining even in the absence of continuing input. After receiving further input, these assemblies create coexisting spiking activity rather than creating competitive spiking activity, which promotes the reactivation and maintenance of the information [46, 70]. The reconstructed and maintained information in the beta oscillations primarily serve the endogenous top-down-controlled processing through long-range connections[18] and are involved in making top-down predictions. Hence, we suggest that in our results, the reactivation of emotional episodes in the beta band contributes to the distinct connectivity patterns for different emotion groups.

In addition, we found that several prediction channels with high prediction accuracy are common in theta and beta bands (fig-5a). The neural activity in brain regions operating in different frequency bands might be contributing to different functionalities. Likewise, a brain stimulation study reported that stimulation of the same brain regions with theta and beta bursts, dissociatively, influenced memory encoding and semantic prediction, respectively [26]. So, the same brain regions can participate in different cognitive abilities at the command of network-specific rhythmic activity. The brain regions we found in our results (including mPFC, MTG, PHG, PCC, IFG, and PC) are reported for episodic and semantic memory processes [62] in theta and beta bands, respectively. Irrespective of whether we remember the past or envisage the future, episodic and semantic elements are inextricably intertwined. Furthermore, the interaction between episodic memory and semantic memory is vital for event construction. The semantic element provides the necessary conceptual or organizational framework for the detailed emotional events to be reconstructed and experienced.

Hence, we suggest that activity patterns in theta and beta bands might be due to the emotional event re-construction process involving the semantic conceptual framework of the already learned emotion concept and representation of the episodic event in this framework. This reconstructed mental event is influencing the cortical processing in a top-down manner to re-experience the situated emotion [12, 91].

### 4.2. Including valence and arousal, other self-assessment emotion features reveal constitutive parts of the emotion mechanism

We observed high prediction accuracy with the channels around the midline and central part of the brain to classify self-assessment emotion features, namely dominance, liking, familiarity, and relevance. On the contrary, valence and arousal are mostly predicted by lateral brain regions (fig-5c). The midline regions and regions around the central part of the brain are reported in self-evaluation [56, 29], autobiographical self [8], mental simulation [28] and event representation [78]. Midline regions comprise “default mode network (DMN)”, which is involved in event construction by integrating current contexts with the internally generated mentalization based on previous experiences to facilitate task performance. The DMN regions play a key role in integrating the acquired current salient knowledge from the stimuli with the internal representations generated using autobiographical and long-term memory to perceive and act on the ongoing interaction with the environment [78].

Familiar and self-relevant contexts can cause the emergence of the emotional state dynamically, even in the absence of external stimuli. For instance, [47] showed that spontaneous spatio-temporal coherence in human resting-state brain activity could be categorized into unique emotional states based on endogenous representations that covary with individual differences in mood, personality traits and self-reported feelings. When a familiar context is encountered, it re-activated event representation as per the memory of the past event. This mental representation can trigger a prediction of what comes next to facilitate responsiveness. For instance, it is shown in pattern classification based fMRI study [44] that given the familiar contextual cue about a novel item, the predictability was higher for the novel item than if the unfamiliar and ambiguous cue context is presented. Furthermore, in a study, we have shown that the context familiarity influences the positive and negative feelings depending on the predictions [53].

In the naturalistic environment, event re-construction brings together information about people, objects, sequences of actions and their consequences to form a multi-modal representation of events. These events models work as generative models to make predictions and guide behaviour [78]. Hence, considering emotion features other than valence and arousal in the emotion experiment paradigm reveals regions involved in the re-construction of emotional events or episodes.

### 4.3. Contribution of DMN in emotion construction

We are getting activity in the medial prefrontal cortex (mPFC), posterior cingulate cortex (PCC)/precuneus(PC), inferior parietal lobe, lateral temporal cortex, and parahippocampal gyrus (fig-6a, 6b), which are reported in the literature as part of the default mode network (DMN) [77]. The DMN simulates mental models of the world from different points of view at different time points. The multi-modal sensorimotor summaries of DMN become more detailed and particularized as they cascade out to primary sensory and motor regions[13].

The DMN is the active sense-making construction network that integrates the salient information in the environment with the interoception, and internal imagery [33, 89]. The information integration and construction process follow a process-memory hierarchy distributed across all cortical areas and dynamically synthesize newly arrived salient input at their preferred timescales [21]. Early sensory areas integrate frequently changing information, whereas association cortices integrate changes over hundreds of milliseconds, and high order areas operate in seconds and overlap with the DMN network. DMN network integrates salient sensory information with intrinsic prior beliefs to form rich contextual information. For instance, a narrative story, with equally possible ambiguous interpretations, is used to test the similar and different interpretations as per the intrinsic beliefs. A stronger aligned activity is reported in the DMN regions for the group with similar interpretation than for the group with the different interpretation [90]. In addition, even if the content is the same, the neural pattern might change for a different interpretation (as the participants in our experiment reported different subjective ratings for the same content; supplementary fig-5).

Constructive theory, one of the advocative theories of involvement of DMN in emotion processing, advocates that a person experiences any emotion when they experience a set of highly situated physiological, behavioural, and contextual features known as conceptualization. Conceptualization is the predictive process that utilizes prior experience and learning to make meaning of the concrete features[12]. The conceptualization extends across feature hierarchy from sensory features, multi-modal sensory information to the higher level of abstraction. DMN fits for the role of the high level of abstraction of emotions given its neuroanatomical structure [22]. Neuroanatomically, DMN has lower neuronal density and higher connection distances compared to sensory areas, which are high in neuronal density and low in connection distance. In addition, DMN nodes are observed to be functionally distant regions from the early sensory cortices [72, 50]. Hence, DMN encodes features at a higher level of abstraction, which is generalized across diverse modalities, conditions & episodes [38].

The re-construction of emotional episodes within the conceptual framework forms the coherent mental representation [62]. DMN plays a central role in this re-construction process as it includes brain regions that are reported to be overlapping with the episodic memory (EM) and semantic memory (SM) related tasks [74, 43]. For instance, the episodic memory task related to recollection or familiarity and semantic memory task related to word concept have overlapping activity in the DMN regions, including PCC, IPL, and mPFC [43]. In addition, the activity related to the episodic task takes place in the theta band, whereas beta-band activity facilitates semantic memory task [26]. Additionally, the theta neurofeedback training increases connectivity strength in the DMN network along with increased emotion awareness and regulation [39]. So, episodic activity in the theta band and semantic activity in the beta band involving DMN regions, we suggest, facilitates the construction of emotional events, which are embodied and experienced through the sensory-motor and visceromotor networks [13].

### 4.4. The role of sensorimotor network and salience network in emotion processing and experience

Emotional Experiences are inherently embodied, and action is closely related to emotion. During development, by mimicking the bodily actions, the child learns about simple emotions from the mother, which later, in adulthood, get enriched by social interactions[81]. These learned bodily experiences and processes are used to infer self emotional states [81, 54, 84] as well as others (by mimicking and embodying others actions) [27, 86]. Simulation can take place without conscious effort and activate the neural substrates for both the recognition and experience of ambiguous as well as prototype emotions[55]. Research with humans as well as with the non-human primates suggest that sensorimotor regions especially, somatomotor and parietal regions play an essential role in imitating actions when observing others’ actions or re-activating learned actions to understand others’ intentions and emotions [75, 63].

Lesion and brain stimulation studies also suggest that sensorimotor regions play an essential role in embodiment and emotional experience. For instance, In a lesion-overlap study, patients with lesions in somatosensory cortices(S1 and S2) and insula, as well as in IFG, motor, and premotor cortices was associated with impairments in emotion prosody perception [3]. rTMS stimulation over S2 with the 1Hz repetitive TMS pulses for 12 mins affected the detection of emotional prosody more than the emotional meaning[3]. In another experiment [85], where participants had to match the presented faces for emotional expressions, stimulation over S1 regions, involved in explicit vs incidental processing of facially expressed emotions, influenced the emotion recognition and matching task performance. The ability of emotion discrimination is disrupted when theta-burst is applied over S1, and premotor cortex [59, 10]. Hence, the sensorimotor regions play a critical role in mimicking others’ actions and re-activating the sensory-motor knowledge to recognize emotional faces, voices, and bodies and in experiencing the perceived emotions.

Another network, we have found in our results, called the salience network or visceromotor network, including mainly ACC & aINS (referred to as “domain-general system”) co-activated in the contexts of diverse tasks and conditions and participated in autonomic function and self-awareness [23, 25]. The domain-general role of salience network is reported in socio-emotional behaviours, including emotions [17], theory of mind, and behavioural mirroring [5]. Furthermore, the dissociative role of aINS and ACC relates its activity with visceromotor autonomic regulation. The frontoinsula (aINS) is reported as a major afferent cortical hub for receiving viscero-autonomic feedback. Complementary to it, ACC activity is responsible for generating autonomic visceral and behavioural responses and is reported to be efferent visceromotor cortical hub [71]. Interaction among these regions responds to homeostatically relevant internal or external stimuli and imbue these stimuli with emotional weight [71], hence, embody the emotions. Irrespective of positive or negative values (i.e., reinforcing or penalizing, respectively), the SN responds to homeostatically relevant stimuli and outcomes [16]. Hence, the salience or visceromotor network integrates autonomic feedback and responses with internal representations and environmental demands [71].

Given the overlapping of high predictability regions observed in our results with the DMN, visceromotor, and sensorimotor network, we suggest that DMN is actively constructing emotional episodes given the salient information, which is then acting on the body and the environment in order to sample the sensory information from the body and environment confirming the mentally reconstructed emotional episodes. These emotional episodes, which are being reconstructed using DMN network and embodied with the help of action-perception cycle in visceromotor and sensorimotor network, are felt and reported as an emotion category by the subject [15, 73, 12].

## 5. Conclusions

In the presented analysis, we observed that the connectivity pattern in the theta and beta band is better able to differentiate different emotion groups than the other bands. The theta-beta interaction performs the emotion event construction considering the past experiences and current context in the conceptual framework of abstract emotion.

In addition, prediction channels are localized to sensorimotor network, salience network, and DMN. These brain regions form distinct emotional representations and hence categorize the emotions. The sampling of salient information and feeling of en-activated and reconstructed emotional episodes engages salient, sensory-motor, and visceromotor networks. So, the emotional episode is embodied and felt in sensorimotor and visceromotor networks (involving somatomotor, Ins, ACC). However, further research modelling the dynamic causality among brain regions and brain stimulation studies for functional specificity is needed to understand the mechanism of emotion in the brain.

We also would like to comment on shifting the paradigm in emotion research towards using more ecologically valid stimuli so that the very close to real-life scenarios can be mimicked in the controlled environment and brain correlates of affect and emotions can be adequately probed. We have reported in this article that self-assessment scales correlated brain activities can be dissected in time and associated mental functionalities to disentangle the contribution of these functionalities in the processing of emotions. Consideration of constituent component dynamics would help in understanding the mechanism of emotion better. In this work, we tried to do some justice in this line. However, this work can be improved further by considering the following points.

1. It is better to collect both the information about space and time using multi-modal neuroimaging methods.
2. Peripheral autonomic activities contain significant emotion-related information. By applying more peripheral sensors on the body, the findings of neural patterns can be cross-verified. In addition, the dynamical association between brain activity and autonomic activity can be used to understand the mutual interactions for emotion processing.
3. Increasing the number of participants will indeed generalize the study for a broader audience.
4. Study on temporal dependence between self-assessment scales can be done.

## Data Availability

Dataset is available at openneuro [51].

## Acknowledgment

We wish to extend our sincere thanks to Prof. Narayanan Srinivasan, CBCS, University of Allahabad. He kindly gave the SM some very useful pieces of advice related to the experiment. Dr Sonia baloni ray, IIIT-Delhi, has introduced SM to the EEG technology and taught him how to use it. She demonstrated it to him by letting him participate in her attention study. Then, we would like to thank Mr Amit Tiwary (then M.tech student), Mr Pravin Srivastav (then PhD candidate), and Ms Anandpreet Kaur (then PhD candidate) for helping us in conducting the validation study.

## References

[1] ,2004. Chapter 46 - nonparametric permutation tests for functional neuroimaging, in: Frackowiak, R.S., Friston, K.J., Frith, C.D., Dolan, R.J., Price, C.J., Zeki, S., Ashburner, J.T., Penny, W.D. (Eds.), Human Brain Function (Second Edition). second edition ed.. Academic Press, Burlington, pp. 887–910. URL: https://www.sciencedirect.com/science/article/pii/B9780122648410500482, doi:https://doi.org/10.1016/B978-012264841-0/50048-2.

[2] Adolphs, R., 2017. How should neuroscience study emotions? by distinguishing emotion states, concepts, and experiences. Social cognitive and affective neuroscience 12, 24–31. doi:10.1093/scan/nsw153.

[3] Adolphs, R., Damasio, H., Tranel, D., 2002. Neural systems for recognition of emotional prosody: a 3-d lesion study. Emotion 2, 23. doi:10.1037/1528-3542.2.1.23.

[4] Adolphs, R., Nummenmaa, L., Todorov, A., Haxby, J.V., 2016. Data-driven approaches in the investigation of social perception. Philosophical Transactions of the Royal Society B: Biological Sciences 371, 20150367. doi:10.1098/rstb.2015.0367.

[5] Alcalá-López, D., Vogeley, K., Binkofski, F., Bzdok, D., 2019. Building blocks of social cognition: mirror, mentalize, share? Cortex 118, 4–18. doi:10.1016/j.cortex.2018.05.006.

[6] Andric, M., Goldin-Meadow, S., Small, S.L., Hasson, U., 2016. Repeated movie viewings produce similar local activity patterns but different network configurations. NeuroImage 142, 613–627. doi:10.1016/j.neuroimage.2016.07.061.

[7] Apps, M.A., Tsakiris, M., 2013. Predictive codes of familiarity and context during the perceptual learning of facial identities. Nature communications 4, 1–10. doi:10.1038/ncomms3698.

[8] Araujo, H.F., Kaplan, J., Damasio, A., 2013. Cortical midline structures and autobiographical-self processes: an activation-likelihood estimation meta-analysis. Frontiers in human neuroscience 7, 548. doi:10.3389/fnhum.2013.00548.

[9] Association, A.P., Association, A.P., et al., 2013. Diagnostic and statistical manual of mental disorders: Dsm-5. United States.

[10] Banissy, M.J., Sauter, D.A., Ward, J., Warren, J.E., Walsh, V., Scott, S.K., 2010. Suppressing sensorimotor activity modulates the discrimination of auditory emotions but not speaker identity. Journal of Neuroscience 30, 13552–13557. doi:10.1523/JNEUROSCI.0786-10.2010.

[11] Bao, F.S., Liu, X., Zhang, C., 2011. Pyeeg: an open source python module for eeg/meg feature extraction. Computational intelligence and neuroscience 2011. doi:10.1155/2011/406391.

[12] Barrett, L.F., 2017. The theory of constructed emotion: an active inference account of interoception and categorization. Social cognitive and affective neuroscience 12, 1–23. doi:10.1093/scan/nsw154.

[13] Barrett, L.F., Quigley, K.S., Hamilton, P., 2016. An active inference theory of allostasis and interoception in depression. Philosophical Transactions of the Royal Society B: Biological Sciences 371, 20160011. doi:10.1098/rstb.2016.0011.

[14] Barrett, L.F., Russell, J.A., 1999. The structure of current affect: Controversies and emerging consensus. Current directions in psychological science 8, 10–14. doi:10.1111/1467-8721.00003.

[15] Barrett, L.F., Simmons, W.K., 2015. Interoceptive predictions in the brain. Nature reviews neuroscience 16, 419–429. doi:10.1038/nrn3950.

[16] Bartra, O., McGuire, J.T., Kable, J.W., 2013. The valuation system: a coordinate-based meta-analysis of bold fmri experiments examining neural correlates of subjective value. Neuroimage 76, 412–427. doi:10.1016/j.neuroimage.2013.02.063.

[17] Bastiaansen, J.A., Thioux, M., Keysers, C., 2009. Evidence for mirror systems in emotions. Philosophical Transactions of the Royal Society B: Biological Sciences 364, 2391–2404. doi:10.1098/rstb.2009.0058.

[18] Bastos, A.M., Vezoli, J., Bosman, C.A., Schoffelen, J.M., Oostenveld, R., Dowdall, J.R., De Weerd, P., Kennedy, H., Fries, P., 2015. Visual areas exert feedforward and feedback influences through distinct frequency channels. Neuron 85, 390–401. doi:10.1016/j.neuron.2014.12.018.

[19] Blascovich, J., Mendes, W.B., 2000. Challenge and threat appraisals: The role of affective cues..

[20] Bradley, M.M., Lang, P.J., 1994. Measuring emotion: the self-assessment manikin and the semantic differential. Journal of behavior therapy and experimental psychiatry 25, 49–59. doi: 10.1016/0005-7916(94)90063-9.

[21] Chang, C.H., Lazaridi, C., Yeshurun, Y., Norman, K.A., Hasson, U., 2021. Relating the past with the present: Information integration and segregation during ongoing narrative processing. Journal of Cognitive Neuroscience 33, 1106–1128.

[22] Collins, C.E., Airey, D.C., Young, N.A., Leitch, D.B., Kaas, J.H., 2010. Neuron densities vary across and within cortical areas in primates. Proceedings of the National Academy of Sciences 107, 15927–15932. doi:10.1073/pnas.1010356107.

[23] Craig, A.D., Craig, A., 2009. How do you feel–now? the anterior insula and human awareness. Nature reviews neuroscience 10.

[24] Critchley, H.D., Garfinkel, S.N., 2017. Interoception and emotion. Current opinion in psychology 17, 7–14. doi:10.1016/j.copsyc.2017.04.020.

[25] Critchley, H.D., Nagai, Y., Gray, M.A., Mathias, C.J., 2011. Dissecting axes of autonomic control in humans: insights from neuroimaging. Autonomic Neuroscience 161, 34–42. doi:10.1016/j.autneu.2010.09.005.

[26] Dave, S., VanHaerents, S., Voss, J.L., 2020. Cerebellar theta and beta noninvasive stimulation rhythms differentially influence episodic memory versus semantic prediction. Journal of Neuroscience 40, 7300–7310. doi:10.1523/JNEUROSCI.0595-20.2020.

[27] Decety, J., Meyer, M., 2008. From emotion resonance to empathic understanding: A social developmental neuroscience account. Development and psychopathology 20, 1053–1080. doi:10.1017/S0954579408000503.

[28] Faul, L., Jacques, P.L.S., DeRosa, J.T., Parikh, N., De Brigard, F., 2020. Differential contribution of anterior and posterior midline regions during mental simulation of counterfactual and perspective shifts in autobiographical memories. NeuroImage 215, 116843. doi:10.1016/j.neuroimage.2020.116843.

[29] Flagan, T., Beer, J.S., 2013. Three ways in which midline regions contribute to self-evaluation. Frontiers in human neuroscience 7, 450. doi: 10.3389/fnhum.2013.00450.

[30] Gianotti, L.R., Faber, P.L., Schuler, M., Pascual-Marqui, R.D., Kochi, K., Lehmann, D., 2008. First valence, then arousal: the temporal dynamics of brain electric activity evoked by emotional stimuli. Brain topography 20, 143–156. doi:10.1007/s10548-007-0041-2.

[31] Goldberg, H., Preminger, S., Malach, R., 2014. The emotion–action link? naturalistic emotional stimuli preferentially activate the human dorsal visual stream. Neuroimage 84, 254–264. doi:10.1016/j.neuroimage.2013.08.032.

[32] Guex, R., Méndez-Bértolo, C., Moratti, S., Strange, B.A., Spinelli, L., Murray, R.J., Sander, D., Seeck, M., Vuilleumier, P., Domínguez-Borràs, J., 2020. Temporal dynamics of amygdala response to emotion-and action-relevance. Scientific Reports 10, 1–16. doi:10.1038/s41598-020-67862-1.

[33] Hassabis, D., Maguire, E.A., 2009. The construction system of the brain. Philosophical Transactions of the Royal Society B: Biological Sciences 364, 1263–1271. doi:10.1098/rstb.2008.0296.

[34] Headley, D.B., Paré, D., 2013. In sync: gamma oscillations and emotional memory. Frontiers in behavioral neuroscience 7, 170. doi:10.3389/fnbeh.2013.00170.

[35] Herbert, C., Sfärlea, A., Blumenthal, T., 2013. Your emotion or mine: labeling feelings alters emotional face perceptionan erp study on automatic and intentional affect labeling. Frontiers in Human Neuroscience 7, 378. doi:10.3389/fnhum.2013.00378.

[36] Hesp, C., Smith, R., Parr, T., Allen, M., Friston, K.J., Ramstead, M.J., 2021. Deeply felt affect: The emergence of valence in deep active inference. Neural Computation 33, 1–49.

[37] Holland, A.C., Kensinger, E.A., 2010. Emotion and autobiographical memory. Physics of life reviews 7, 88–131.

[38] Huntenburg, J.M., Bazin, P.L., Margulies, D.S., 2018. Large-scale gradients in human cortical organization. Trends in cognitive sciences 22, 21–31. doi:10.1016/j.tics.2017.11.002.

[39] Imperatori, C., Della Marca, G., Amoroso, N., Maestoso, G., Valenti, E.M., Massullo, C., Carbone, G.A., Contardi, A., Farina, B., 2017. Alpha/theta neurofeedback increases mentalization and default mode network connectivity in a non-clinical sample. Brain topography 30, 822–831. doi:10.1007/s10548-017-0593-8.

[40] Jerram, M., Lee, A., Negreira, A., Gansler, D., 2014. The neural correlates of the dominance dimension of emotion. Psychiatry Research: Neuroimaging 221, 135–141. doi:10.1016/j.pscychresns.2013.11.007.

[41] Jimenez, K.B., Abdelgabar, A.R., De Angelis, L., McKay, L.S., Keysers, C., Gazzola, V., 2020. Changes in brain activity following the voluntary control of empathy. Neuroimage 216, 116529.

[42] Jolly, E., Chang, L., 2021. Multivariate spatial feature selection in fmri. Social Cognitive and Affective Neuroscience.

[43] Kim, H., 2016. Default network activation during episodic and semantic memory retrieval: a selective meta-analytic comparison. Neuropsychologia 80, 35–46. doi:10.1016/j.neuropsychologia.2015.11.006.

[44] Kim, H., Schlichting, M.L., Preston, A.R., Lewis-Peacock, J.A., 2020. Predictability changes what we remember in familiar temporal contexts. Journal of cognitive neuroscience 32, 124–140. doi:10.1162/jocn_a_01473.

[45] Koelstra, S., Muhl, C., Soleymani, M., Lee, J.S., Yazdani, A., Ebrahimi, T., Pun, T., Nijholt, A., Patras, I., 2011. Deap: A database for emotion analysis; using physiological signals. IEEE transactions on affective computing 3, 18–31. doi:10.1109/T-AFFC.2011.15.

[46] Kopell, N., Whittington, M.A., Kramer, M.A., 2011. Neuronal assembly dynamics in the beta1 frequency range permits short-term memory. Proceedings of the National Academy of Sciences 108, 3779–3784. doi:10.1073/pnas.1019676108.

[47] Kragel, P.A., Knodt, A.R., Hariri, A.R., LaBar, K.S., 2016. Decoding spontaneous emotional states in the human brain. PLoS biology 14, e2000106. doi:10.1371/journal.pbio.2000106.

[48] Landman, L.L., van Steenbergen, H., 2020. Emotion and conflict adaptation: the role of phasic arousal and self-relevance. Cognition and Emotion 34, 1083–1096.

[49] Lowe, R., 2011. The feeling of action tendencies: on the emotional regulation of goal-directed behavior. Frontiers in psychology 2, 346. doi: 10.3389/fpsyg.2011.00346.

[50] Margulies, D.S., Ghosh, S.S., Goulas, A., Falkiewicz, M., Huntenburg, J.M., Langs, G., Bezgin, G., Eickhoff, S.B., Castellanos, F.X., Petrides, M., et al., 2016. Situating the default-mode network along a principal gradient of macroscale cortical organization. Proceedings of the National Academy of Sciences 113, 12574–12579. doi:10.1073/pnas.1608282113.

[51] Mishra, S., Asif, M., Tiwary, U.S., 2021a. “dataset on emotion with naturalistic stimuli”. doi:10.18112/openneuro.ds003751.v1.0.0.

[52] Mishra, S., Asif, M., Tiwary, U.S., 2021b. Dataset on emotions using naturalistic stimuli (dens). bioRxiv URL: https://www.biorxiv.org/content/early/2021/08/05/2021.08.04.455041, doi:10.1101/2021.08.04.455041, arXiv:https://www.biorxiv.org/content/early/2021/08/05/2021.08.04.455041.full.pdf.

[53] Mishra, S., Tiwary, U.S., 2021. The degree of context un/familiarity impacts the emotional feeling and preaware cardiac-brain activity: a study with emotionally salient naturalistic paradigm using dens dataset. bioRxiv URL: https://www.biorxiv.org/content/early/2021/08/08/2021.08.07.455496, doi:10.1101/2021.08.07.455496, arXiv:https://www.biorxiv.org/content/early/2021/08/08/2021.08.07.455496.full.pdf.

[54] Niedenthal, P.M., Mermillod, M., Maringer, M., Hess, U., 2010. The simulation of smiles (sims) model: Embodied simulation and the meaning of facial expression. Behavioral and brain sciences 33, 417. doi:10.1017/S0140525X10000865.

[55] Niedenthal, P.M., Ric, F., 2017. Psychology of emotion. Psychology Press. doi:10.4324/9781315276229.

[56] Northoff, G., Bermpohl, F., 2004. Cortical midline structures and the self. Trends in cognitive sciences 8, 102–107. doi:10.1016/j.tics.2004.01.004.

[57] Nummenmaa, L., Glerean, E., Viinikainen, M., Jääskeläinen, I.P., Hari, R., Sams, M., 2012. Emotions promote social interaction by synchronizing brain activity across individuals. Proceedings of the National Academy of Sciences 109, 9599–9604. doi:10.1073/pnas.1206095109.

[58] Nummenmaa, L., Saarimäki, H., Glerean, E., Gotsopoulos, A., Jääskeläinen, I.P., Hari, R., Sams, M., 2014. Emotional speech synchronizes brains across listeners and engages large-scale dynamic brain networks. NeuroImage 102, 498–509. doi:10.1016/j.neuroimage.2014.07.063.

[59] Pitcher, D., Garrido, L., Walsh, V., Duchaine, B.C., 2008. Transcranial magnetic stimulation disrupts the perception and embodiment of facial expressions. Journal of Neuroscience 28, 8929–8933. doi:10.1523/JNEUROSCI.1450-08.2008.

[60] Raichle, M.E., 2015. The brain’s default mode network. Annual review of neuroscience 38, 433–447. doi:10.1146/annurev-neuro-071013-014030.

[61] Rameson, L.T., Satpute, A.B., Lieberman, M.D., 2010. The neural correlates of implicit and explicit self-relevant processing. NeuroImage 50, 701–708. doi:10.1016/j.neuroimage.2009.12.098.

[62] Renoult, L., Irish, M., Moscovitch, M., Rugg, M.D., 2019. From knowing to remembering: the semantic–episodic distinction. Trends in Cognitive Sciences 23, 1041–1057. doi:10.1016/j.tics.2019.09.008.

[63] Ross, P., Atkinson, A.P., 2020. Expanding simulation models of emotional understanding: the case for different modalities, body-state simulation prominence, and developmental trajectories. Frontiers in psychology 11, 309. doi:10.3389/fpsyg.2020.00309.

[64] Saarimäki, H., 2021. Naturalistic stimuli in affective neuroimaging: a review.

[65] Sakaki, M., Niki, K., Mather, M., 2012. Beyond arousal and valence: The importance of the biological versus social relevance of emotional stimuli. Cognitive, Affective, & Behavioral Neuroscience 12, 115–139. doi:10.3758/s13415-011-0062-x.

[66] Sander, D., Grandjean, D., Scherer, K.R., 2018. An appraisal-driven componential approach to the emotional brain. Emotion Review 10, 219–231.

[67] Satpute, A.B., Lindquist, K.A., 2019. The default mode networks role in discrete emotion. Trends in cognitive sciences 23, 851–864.

[68] Satpute, A.B., Nook, E.C., Narayanan, S., Shu, J., Weber, J., Ochsner, K.N., 2016. Emotions in black and white or shades of gray? how we think about emotion shapes our perception and neural representation of emotion. Psychological science 27, 1428–1442.

[69] Scherer, K.R., 2009. The dynamic architecture of emotion: Evidence for the component process model. Cognition and emotion 23, 1307–1351. doi:10.1080/02699930902928969.

[70] Scholz, S., Schneider, S.L., Rose, M., 2017. Differential effects of ongoing eeg beta and theta power on memory formation. PloS one 12, e0171913. doi:10.1371/journal.pone.0171913.

[71] Seeley, W.W., 2019. The salience network: a neural system for perceiving and responding to homeostatic demands. Journal of Neuroscience 39, 9878–9882. doi:10.1523/JNEUROSCI.1138-17.2019.

[72] Sepulcre, J., Sabuncu, M.R., Yeo, T.B., Liu, H., Johnson, K.A., 2012. Stepwise connectivity of the modal cortex reveals the multimodal organization of the human brain. Journal of Neuroscience 32, 10649–10661. doi:10.1523/JNEUROSCI.0759-12.2012.

[73] Seth, A.K., Friston, K.J., 2016. Active interoceptive inference and the emotional brain. Philosophical Transactions of the Royal Society B: Biological Sciences 371, 20160007. doi:10.1098/rstb.2016.0007.

[74] Shapira-Lichter, I., Oren, N., Jacob, Y., Gruberger, M., Hendler, T., 2013. Portraying the unique contribution of the default mode network to internally driven mnemonic processes. Proceedings of the National Academy of Sciences 110, 4950–4955. doi:10.1073/pnas.1209888110.

[75] Spaulding, S., 2013. Mirror neurons and social cognition. Mind & Language 28, 233–257. doi:10.1111/mila.12017.

[76] Spitzer, B., Haegens, S., 2017. Beyond the status quo: a role for beta oscillations in endogenous content (re) activation. eneuro 4. doi:10.1523/ENEURO.0170-17.2017.

[77] Spreng, R.N., Stevens, W.D., Chamberlain, J.P., Gilmore, A.W., Schacter, D.L., 2010. Default network activity, coupled with the frontoparietal control network, supports goal-directed cognition. Neuroimage 53, 303–317. doi:10.1016/j.neuroimage.2010.06.016.

[78] Stawarczyk, D., Bezdek, M.A., Zacks, J.M., 2021. Event representations and predictive processing: The role of the midline default network core. Topics in Cognitive Science 13, 164–186. doi:10.1111/tops.12450.

[79] Van Den Bosch, I., Salimpoor, V., Zatorre, R.J., 2013. Familiarity mediates the relationship between emotional arousal and pleasure during music listening. Frontiers in human neuroscience 7, 534. doi:10.3389/fnhum.2013.00534.

[80] Verma, G.K., Tiwary, U.S., 2014. Multimodal fusion framework: A multiresolution approach for emotion classification and recognition from physiological signals. NeuroImage 102, 162–172. doi:10.1016/j.neuroimage.2013.11.007.

[81] Williams, J.H., Huggins, C.F., Zupan, B., Willis, M., Van Rheenen, T.E., Sato, W., Palermo, R., Ortner, C., Krippl, M., Kret, M., et al., 2020. A sensorimotor control framework for understanding emotional communication and regulation. Neuroscience & Biobehavioral Reviews 112, 503–518. doi:10.1016/j.neubiorev.2020.02.014.

[82] Wilson-Mendenhall, C.D., Barrett, L.F., Simmons, W.K., Barsalou, L.W., 2011. Grounding emotion in situated conceptualization. Neuropsychologia 49, 1105–1127. doi:10.1016/j.neuropsychologia.2010.12.032.

[83] Winecoff, A., Clithero, J.A., Carter, R.M., Bergman, S.R., Wang, L., Huettel, S.A., 2013. Ventromedial prefrontal cortex encodes emotional value. Journal of Neuroscience 33, 11032–11039. doi:10.1523/JNEUROSCI.4317-12.2013.

[84] Winkielman, P., 2010. Embodied and disembodied processing of emotional expressions: insights from autism spectrum disorders. Behavioral and Brain Sciences 33, 463. doi:10.1017/S0140525X10001640.

[85] Winston, J.S., O’doherty, J., Dolan, R.J., 2003. Common and distinct neural responses during direct and incidental processing of multiple facial emotions. Neuroimage 20, 84–97. doi:10.1016/S1053-8119(03)00303-3.

[86] Wood, A., Rychlowska, M., Korb, S., Niedenthal, P., 2016. Fashioning the face: sensorimotor simulation contributes to facial expression recognition. Trends in cognitive sciences 20, 227–240. doi:10.1016/j.tics.2015.12.010.

[87] Yang, K., Tong, L., Shu, J., Zhuang, N., Yan, B., Zeng, Y., 2020. High gamma band eeg closely related to emotion: evidence from functional network. Frontiers in human neuroscience 14, 89. doi:10.3389/fnhum.2020.00089.

[88] Yarkoni, T., Poldrack, R.A., Nichols, T.E., Van Essen, D.C., Wager, T.D., 2011. Large-scale automated synthesis of human functional neuroimaging data. Nature methods 8, 665–670. doi:10.1038/nmeth.1635.

[89] Yeshurun, Y., Nguyen, M., Hasson, U., 2021. The default mode network: where the idiosyncratic self meets the shared social world. Nature Reviews Neuroscience 22, 181–192.

[90] Yeshurun, Y., Swanson, S., Simony, E., Chen, J., Lazaridi, C., Honey, C.J., Hasson, U., 2017. Same story, different story: the neural representation of interpretive frameworks. Psychological science 28, 307–319. doi:10.1177/0956797616682029.

[91] Zhou, P., Critchley, H., Garfinkel, S., Gao, Y., 2021. The conceptualization of emotions across cultures: A model based on interoceptive neuroscience. Neuroscience & Biobehavioral Reviews.

