## supplementary fig-1, supplementary fig-2, supplementary fig-3, supplementary fig-4, supplementary fig-5, supplementary fig-6, supplementary fig-7 for "Naturalistic paradigm reveals multi-component emotion dynamics in theta and beta bands using DENS dataset"

### 1. SI: EEG Features

I. **Detrended fluctuation analysis(DFA):** The dfa in nutshell is a technique to test the scale free structure in the signal. It is comprised of following steps:
1. The signal $x_1, x_2, ...., x_n$ is integrated to into a new series y = [y(1), y(2),... y(N)] where $y(k) = \Sigma_{i=1}^{k}(x_i - \overline{x})$. $\overline{x}$ is the average of original signal X.
2. The integrated series is then first divided into boxes of equal size n. For each box least-square fitted line is calculated, representing the trend in that box. The process of subtracting this least square line from the integrate series is called detrending.
3. Using the formula $F(n) = \sqrt{\frac{1}{N}\Sigma_{k=1}^{N}[y(k) - y_n(k)]^2}$ the root mean square fluctuation of the integrated series is calculated.
4. The DFA values is the slope of log F(n) to log n.

II. **Power Spectral Intensity(PSI) and Relative Intensity Ratio(RIR):** The original signal is frequency transformed and discrete frequency bands are partitioned into unequal k bins. The power spectral intensity for $k^{th}$ bin is evaluated as:

$$PSI_k = \Sigma_{i=\lfloor N(\frac{f_k}{f_s})\rfloor}^{\lfloor N(\frac{f_{k+1}}{f_s})\rfloor} |X|,\ k = 1, 2, ...., K - 1$$

where, $f_s$ = sampling rate, N = length of the series.

$$RIR_k = \frac{PSI_k}{\Sigma_{k=1}^{K-1} PSI_k}, j = 1, 2, ....., K - 1$$

III. **Spectral entropy:**

$$H = -\frac{1}{log(K)}\Sigma_{i=1}^{K} RIR_i log RIR_i$$

IV. **Petrosian Fractal Dimension(PFD):** PFD for a time-series is defined as:

$$PFD = \frac{log_{10}N}{log_{10}N + log_{10}(\frac{N}{(N+0.4N_\delta)})}$$

where N = series length; $N_\delta$ = number of sign changes in the signal derivative.

V. **Higuchi Fractal Dimension (HFD):** Higuchi fractal dimension is the approximation of the value for the box-counting dimension of the graph of a real-value time-series. The higuchi fractal dimension of a time series for $k \in \{1, ...., k_{max}\}$ and $m \in \{1, ...., k\}$ is the slope of the best-fitting linear function through the data points $\{(log\frac{1}{k}, log L(k))\}$. Where,

$$L(k) = \frac{1}{k}\Sigma_{m=1}^{k} L_m(k)$$

, and

$$L_m(k) = \frac{N-1}{[\frac{N-m}{k}]k^2}\Sigma_{i=1}^{[\frac{N-m}{k}]}|X_N(m + ik) - X_N(m + (i-1)k|$$

VI. **Hjorth mobility and complexity:** Hjorth parameters are indicators of statistical properties used to analyze activity, mobility and complexity of time series in signal processing. We have used the mobility and complexity paraters which are as follows:

$$Mobility = \sqrt{\frac{var(\frac{d_y(t)}{dt})}{var(y(t))}}$$

$$Complexity = \frac{Mobility(\frac{d_y(t)}{dt})}{Mobility(y(t))}$$

where, y(t) is the signal. Mobility represents the mean frequency or the proportion of standard deviation of the power spectrum whereas the complexity represents the change in frequency.

VII. **SVD Entropy:** Dissecting the terms to define SVD and entropy individually. The SVD weights indicate how much of the data set is explained by each vector. Entropy delivers the maximally-noncommittal data set at a given signal-noise ratio, that is to say, the most information with the least artefact.

So the SVD entropy is an indicator of how many vectors are needed for an adequate explanation of the data set. You could say it measures feature-richness in the sense that the higher the entropy of the set of SVD weights, the more orthogonal vectors are required to adequately explain it.

The SVD entropy is defined as

$$H_{SVD} = \Sigma_{i=1}^{M} \overline{\sigma_i} log_2 \overline{\sigma_i}$$

, where M is the number of singular values and $\overline{\sigma_1}, ....., \overline{\sigma_M}$ are normalized singular values such that $\overline{\sigma_i} = \frac{\sigma_i}{\Sigma_{j=1}^{M} \sigma_j}$.

VIII. **Fisher information:** The Fisher information is a way of measuring the amount of information that an observable random variable X carries about an unknown parameter $\theta$ of a distribution that models X. It is defined as:

$$I = \Sigma_{i=1}^{M-1} \frac{(\overline{\sigma_{i+1}} - \overline{\sigma_i})^2}{\overline{\sigma_i}}$$

IX. **Approximate entropy(ApEn):** This entropy measure is used to quantify the amount of regularity and the unpredictability of fluctuations over time-series data.

X. **Hurst Exponent:** The hurst exponent is a measure to quantify a long-term memory of a time series. To calculate the hurst exponent of a time series, first, the accumulated deviation from the mean of time series within range T has to be calculated as follows:

$$X(t,T) = \Sigma_{i=1}^{t} (x_i - \overline{x})$$

where

$$\overline{x} = \frac{1}{T} \Sigma_{i=1}^{T} x_i,\ t \in [1..N]$$

$$\frac{R(T)}{S(t)} = \frac{max(X(t,T)) - min(X(t,T))}{\sqrt{(\frac{1}{T}) \Sigma_{i=1}^{T} [x(t) - \overline{x}]^2}}$$

Then, the hurst exponent is slope of the line produced by plotting $log \frac{R(T)}{S(T)}$ versus $log(T)$

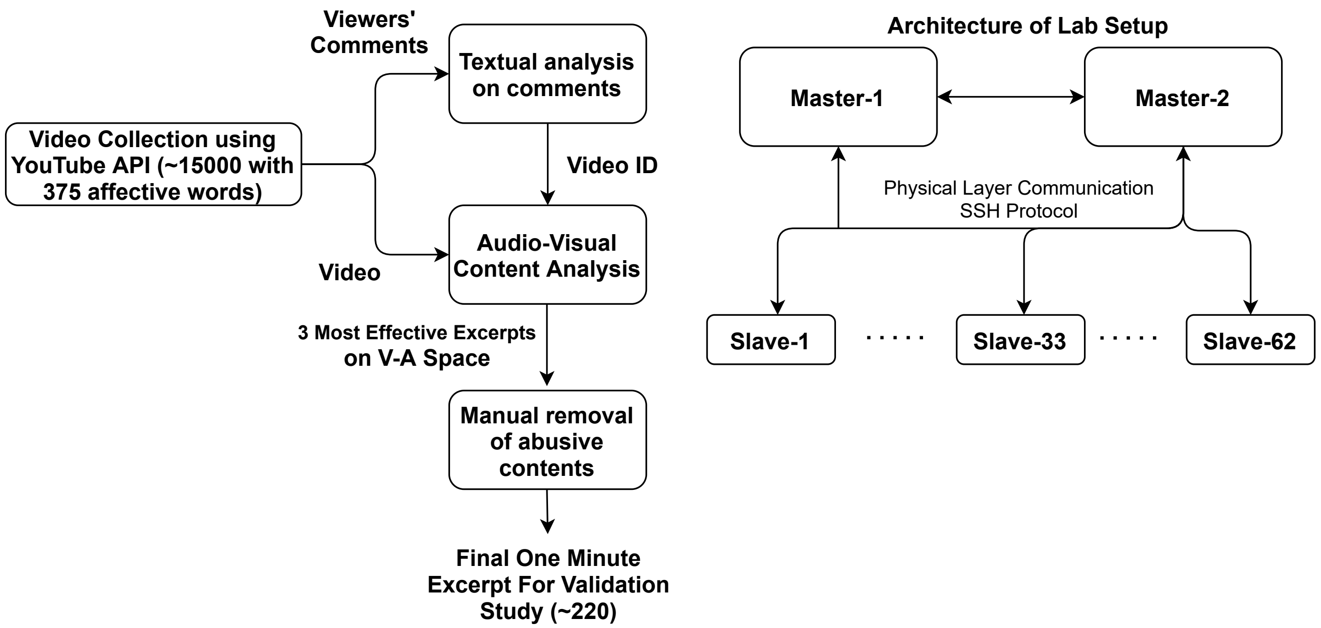


**Figure 1: SI: Stimulus Selection and Validation** Left figure represents the procedure which is used to collect the stimuli from internet and then performing textual and multimedia content analysis to finally select around 220 stimuli. The mulitmedia feature which are calculated on these stimuli are described in [2]. Right figure is representing lab configuration where stimuli validation is carried out. All the system had linux mint operating system installed in it. A master-slave architecture based on ssh protocol is implemented. All the stimuli are are divided into 22 blocks randomly into the master system and then distributed to all the slaves dynamically while students are being informed about the experiment by a slide show presentation. Once, students are clear about instructions they are assigned one of the slave system. To be noted, the distance between participants were around 2 meters so that they are more focused on their own screen (they are also requested for the same). Lights were off and windows were properly shielded from light so that an immersive environment can be created. Three volunteers were monitoring the activities and were also available for any help to the participants. Once the experiment is done, in master system a script run by the experimenter which collected all the responses from the slave system to the master system.

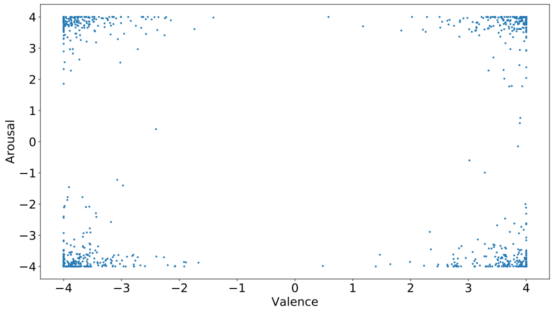


**Figure 2: Multimedia content based valence-arousal plot:** The valence and arousal is calculated using multimedia content analysis and only those one-minute excerpts are selected and plotted here which are on the periphery. The magnitude of vector is calculated using the formula $e_i = \sqrt{a_i^2 + v_i^2}$. Smaller $e_i$ is closer to the neutral state. Excerpts with a higher $e_i$ score are selected for the validation study.

### References


[1] Koelstra, S., Muhl, C., Soleymani, M., Lee, J.S., Yazdani, A., Ebrahimi, T., Pun, T., Nijholt, A., Patras, I., 2011. Deap: A database for emotion analysis; using physiological signals. IEEE transactions on affective computing 3, 18–31. doi:10.1109/T-AFFC.2011.15.

[2] Mishra, S., Asif, M., Tiwary, U.S., 2021. Dataset on emotions using naturalistic stimuli (dens). bioRxiv URL: https://www.biorxiv.org/content/early/2021/08/05/2021.08.04.455041, doi:10.1101/2021.08.04.455041, arXiv:https://www.biorxiv.org/content/early/2021/08/05/2021.08.04.455041.full.pdf.

[3] Warriner, A.B., Kuperman, V., Brysbaert, M., 2013. Norms of valence, arousal, and dominance for 13,915 english lemmas. Behavior research methods 45, 1191–1207. doi:10.3758/s13428-012-0314-x.

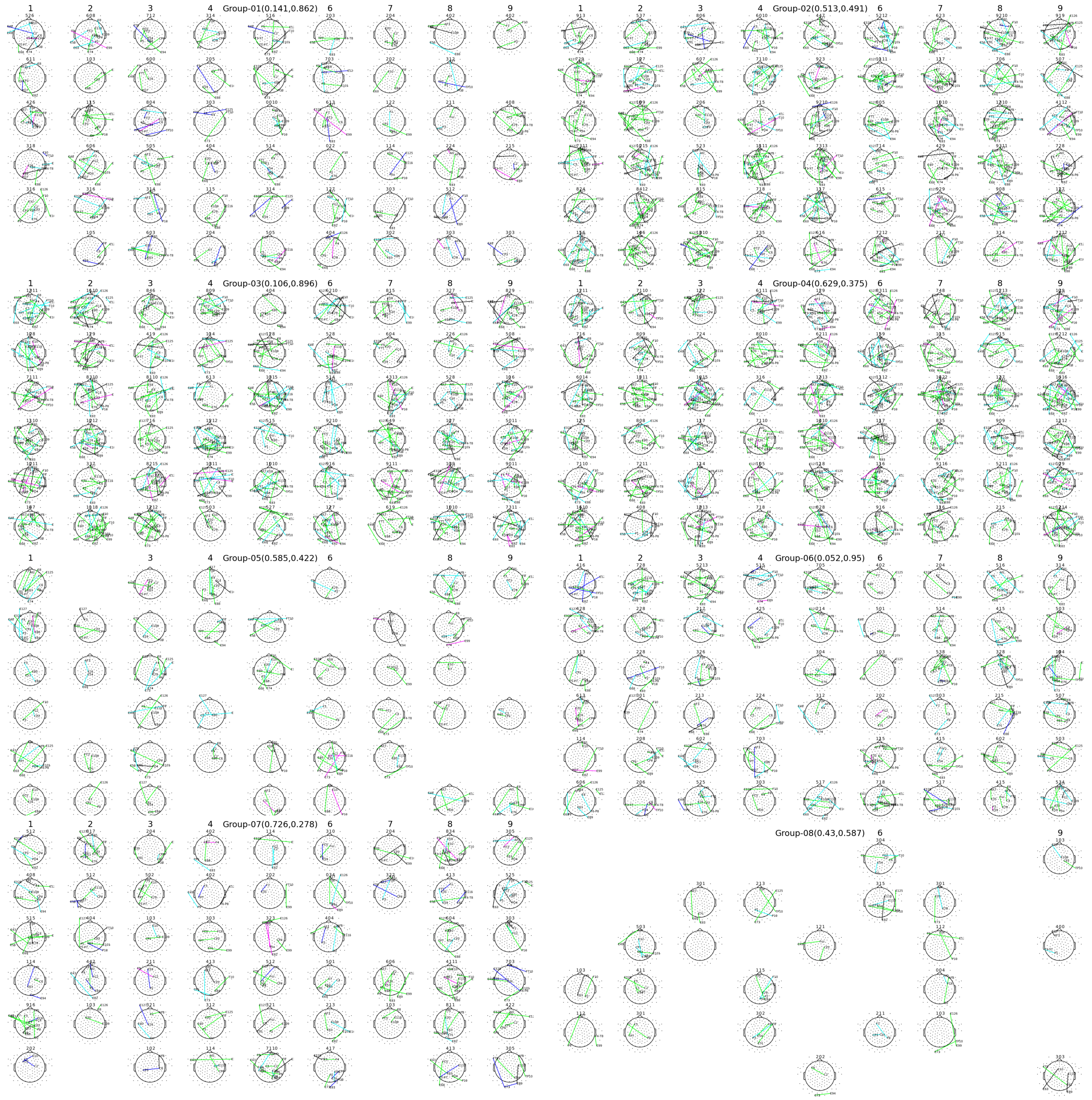

**Figure 3: SI: Connectivity** Using phase locking value (PLV) the connectivity among pair of electrodes is calculated. The connectivity matrix for each emotion is then contrasted with the resting state connectivity matrix. The plots here are showing statistically significant connections calculated using non-parametric permutation test (p-value adjusted for multiple comparison). Row wise top to bottom, delta, theta, alpha, lower beta, upper beta and gamma bands. Column wise left to right segment 1 to segment-9. The blank space are the cell with no significant connections.

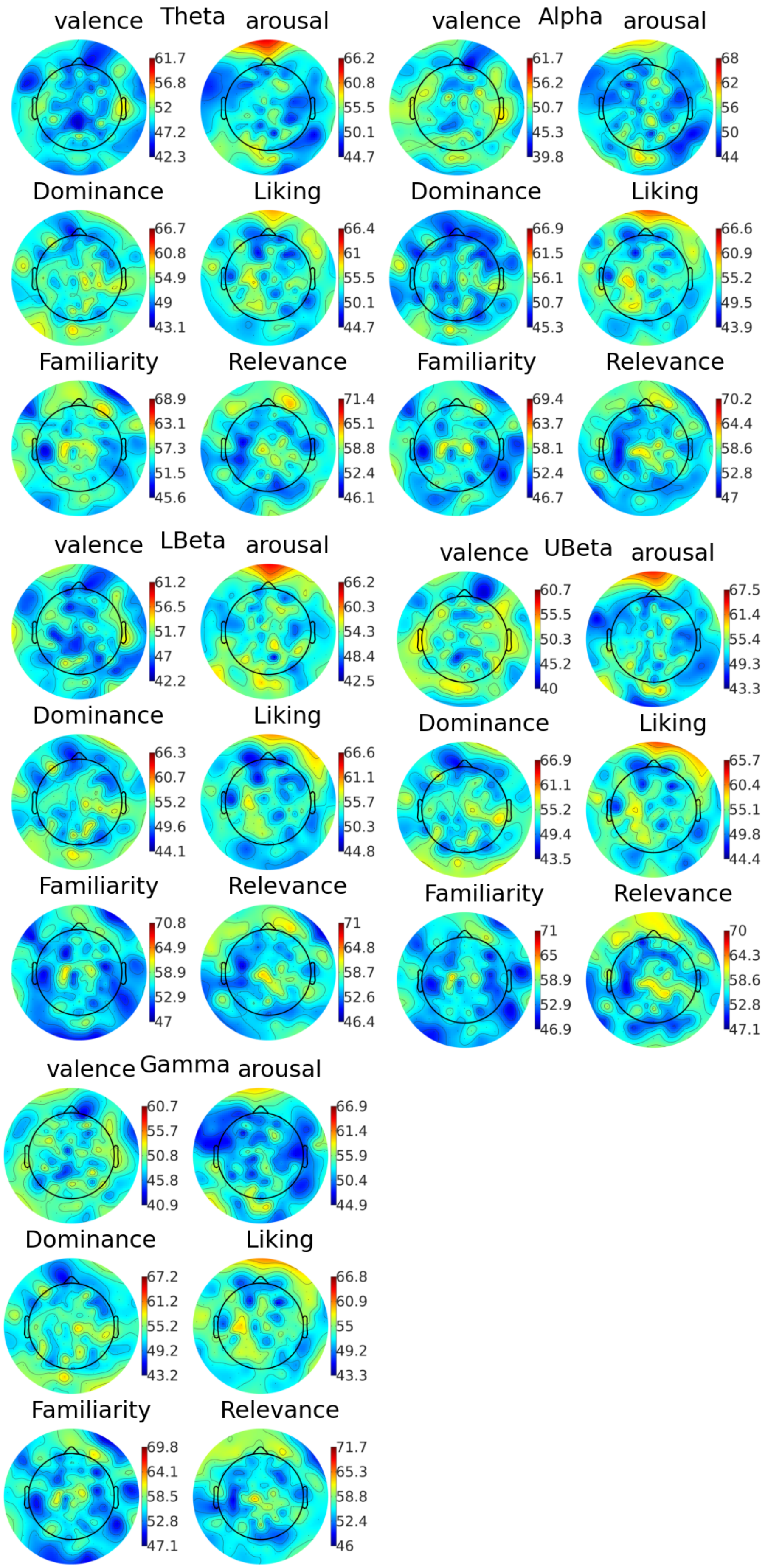




**Figure 4: SI: Prediction Topoplots**

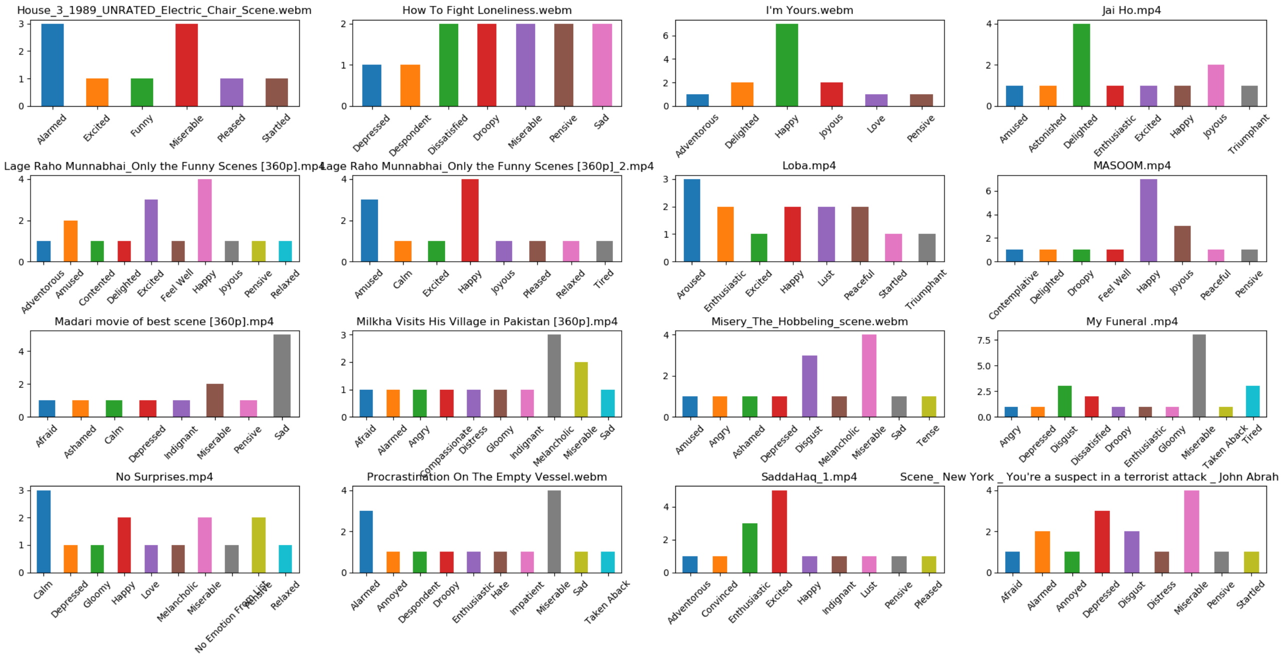


**Figure 5: SI: Emotion Tag Distribution** For each stimuli subjects had responded on the valence and arousal scale(along with the other scales). We took these ratings for each participants and mapped them on the valence-arousal plot for well known previous datasets on emotions, namely, DEAP data [1] and Warinner data [3]. We calculated the domain of an emotion, for both the datasets, by considering mean as the center and standard deviation as the diameter. And, assigned this emotion to all the observations which are inside this circle. We performed this calculation so that it can be shown that how the same stimuli could be conceptualized differently by different participants.

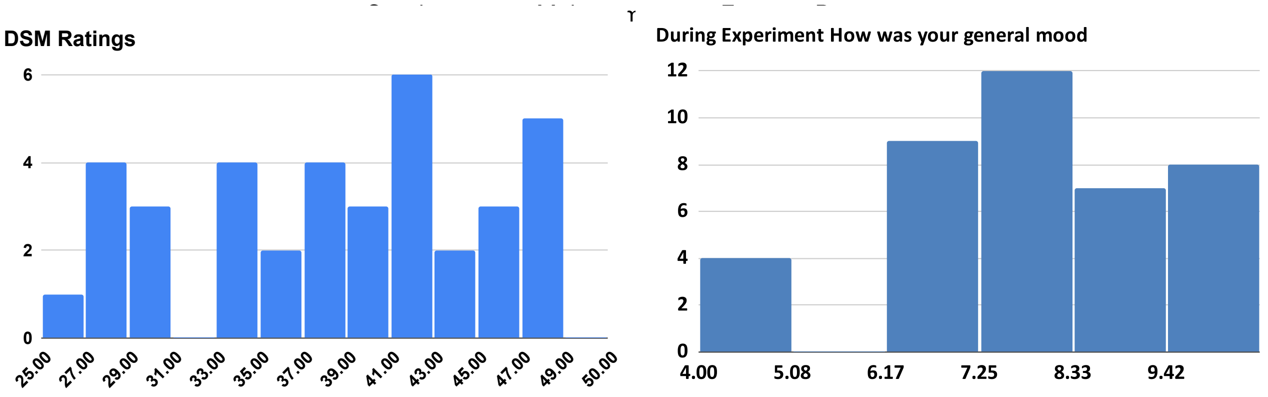


(a) DSM Ratings

(b) General Mood of participants during experiment

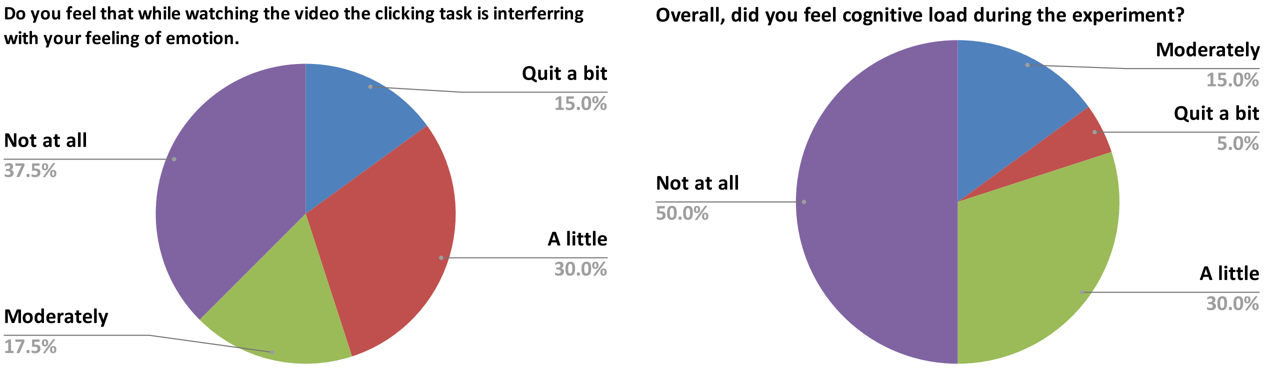


(c) Click is Interfering with emotion feeling

(d) Cognitive Load during experiment

**Figure 6: SI: Participants Questionnaire** (a) Histogram of DSM ratings, (b) overall general mood of the participants while performing the experiment, (c) how much a non-emotional clicking task was interfering with the feeling of emotions, (d) how much cognitive load participants felt during the experiment.

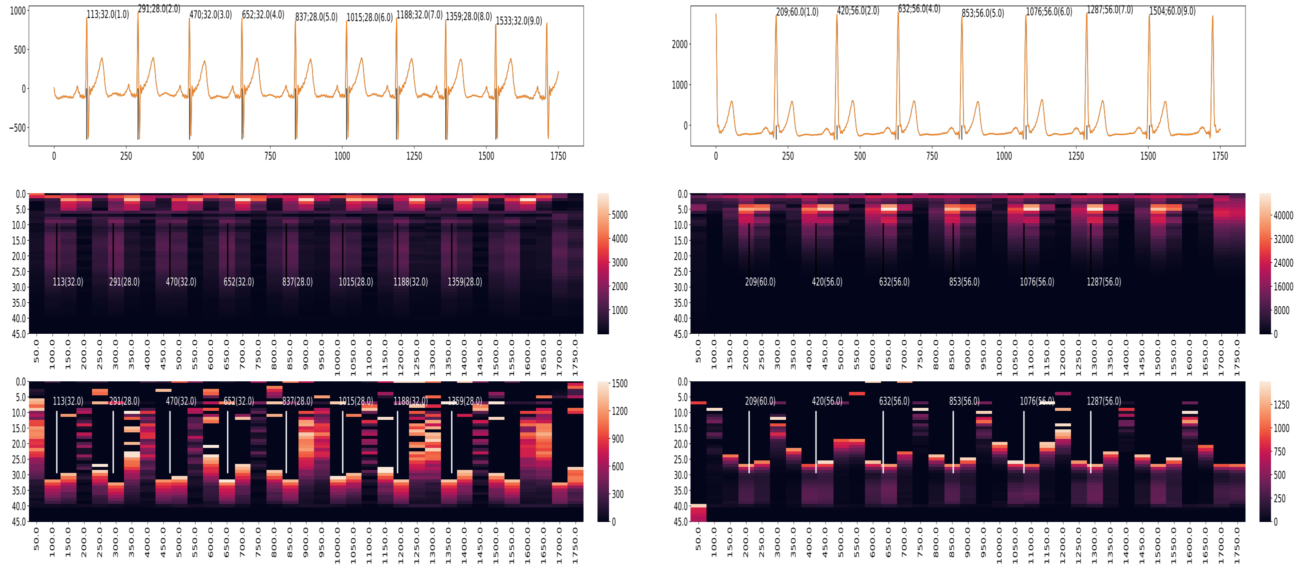

(a)

(b)

**Figure 7: SI:ECG** (a-b) ECG with spectrogram calculated using stft. First diagram in (a-b) is ECG graph, second diagram is spectrogram calculated using stft, and third diagram is emphasizing low magnitude frequencies ($mag > 450$). The annotation in ECG graph is time of Rpeak:duration of Rpeak(segment number). For low R-width, the magnitude of higher frequencies is relatively more than the higher frequency magnitude for high R-width (as shown in the graph relatively more magnitude for 15Hz to 30 Hz for low R-width).
